# Variation in Human Herpesvirus 6B telomeric integration, excision and transmission between tissues and individuals

**DOI:** 10.1101/2021.06.07.447326

**Authors:** M.L. Wood, C. Veal, R. Neumann, N.M. Suárez, J. Nichols, A.J. Parker, D. Martin, S.P.R Romaine, V. Codd, N.J. Samani, A.A. Voors, M. Tomaszewski, L. Flamand, A.J. Davison, N.J. Royle

## Abstract

Human herpesviruses 6A and 6B (HHV-6A/6B) are ubiquitous pathogens that persist lifelong in latent form and can cause severe conditions upon reactivation. They are spread by community-acquired infection of free virus (acqHHV6A/6B) and by germline transmission of inherited chromosomally-integrated HHV-6A/6B (iciHHV-6A/6B) in telomeres. We exploited a hypervariable region of the HHV- 6B genome to investigate the relationship between acquired and inherited virus and revealed predominantly maternal transmission of acqHHV-6B in families. Remarkably, we demonstrate that some copies of acqHHV-6B in saliva from healthy adults gained a telomere, indicative of integration and latency, and that the frequency of viral genome excision from telomeres in iciHHV-6B carriers is surprisingly high and varies between tissues. In addition, newly formed short telomeres generated by partial viral genome release are frequently lengthened, particularly in telomerase-expressing pluripotent cells. Consequently, iciHHV-6B carriers are mosaic for different iciHHV-6B structures, including circular extra-chromosomal forms that have the potential to reactivate. Finally, we show transmission of an HHV-6B strain from an iciHHV-6B mother to her non-iciHHV-6B son. Altogether we demonstrate that iciHHV-6B can readily transition between telomere-integrated and free virus forms.

## Introduction

Human herpesviruses 6A and 6B (HHV-6A/6B; species *Human betaherpesvirus 6A* and *Human betaherpesvirus 6B*) are closely related viruses, sharing approximately 90% sequence identity (Ablashi *et al*, 2014). Their genomes comprise a unique region (U) of approximately 140 kb flanked on each side by a direct repeat (DR) of approximately 8 kb (DRL on the left and DRR on the right). A tandem repeat reiteration is located near each end of DR. T1 at the left end consists of 1-8 kb of telomere (TTAGGG) and degenerate telomere-like repeats (Huang *et al*, 2014; Lindquester & Pellett, 1991; Zhang *et al*, 2017). T2 at the right end is much shorter and contains only (TTAGGG) repeats (Achour *et al*, 2009; Zhang *et al*., 2017). HHV-6A/6B have the capacity to integrate into human telomeres, most likely through a homology-dependent recombination mechanism facilitated by the perfect telomere repeats in T2 (Arbuckle *et al*, 2010; Wallaschek *et al*, 2016b). The precise integration mechanism has not been defined, and searches for regulatory viral or host factors are underway (Gilbert-Girard *et al*, 2017; Gilbert-Girard *et al*, 2020; Wallaschek *et al*, 2016a; Wight *et al*, 2018).

HHV-6A/6B infect over 90% of the global population in early childhood, this usually manifests with mild symptoms but nevertheless often requires medical attention (Asano *et al*, 1994; Hall *et al*, 1994; Kondo *et al*, 1993; Ward & Gray, 1994; Zerr *et al*, 2005). As with most herpesviruses, HHV-6A/6B enter lifelong latency following primary infection, and the most serious impacts on health appear to occur when the virus reactivates. Reactivation of free virus acquired in the community (acqHHV-6A/6B) has been shown to cause viral encephalitis, drug-induced hypersensitivity syndrome/drug reaction with eosinophilia and systemic symptoms (DIHS/DRESS) and acute graft-versus-host disease (aGVHD) (Aihara *et al*, 2003; Eshki *et al*, 2009; Pritchett *et al*, 2012; Prusty *et al*, 2018; Yao *et al*, 2010; Yoshikawa *et al*, 2006). In addition, HHV-6A infection has been associated with multiple sclerosis and other chronic neurological conditions (Yao *et al*., 2010). Herpesviruses typically achieve latency by forming circular DNA episomes within specific cell types, but HHV-6A/6B episomes have not been detected and it has been proposed that latency of these viruses is achieved through telomeric integration (Arbuckle *et al*., 2010).

As a result of historical germline integrations into telomeres, approximately 1% of the population carries inherited chromosomally integrated HHV-6A/6B (iciHHV-6A/6B), usually with a full-length copy of the viral genome in every cell (Arbuckle *et al*., 2010; Daibata *et al*, 1999; Huang *et al*., 2014; Leong *et al*, 2007; Morris *et al*, 1999; Tanaka-Taya *et al*, 2004). HHV-6A/6B appear to have the potential to integrate into any telomere however, most contemporary instances of iciHHV-6A/6B are derived from a relatively small number of ancient integrations in a limited number of chromosome ends (Aswad *et al*, 2021; Greninger *et al*, 2018; Kawamura *et al*, 2017; Liu *et al*, 2018; Liu *et al*, 2020; Miura *et al*, 2018; Tweedy *et al*, 2016; Zhang *et al*., 2017). For example, iciHHV-6B is commonly found integrated into the telomere on the short arm of chromosome 17 (17p) in populations of European descent and iciHHV-6A is commonly found in the long arm of chromosome 22 (22q) in Japan. Integration sites have been determined in a variety of studies by one or a combination of the following methods: Fluorescent *in situ* hybridisation (FISH); PCR amplification from subterminal sequences adjacent to human telomeres (subtelomere) into the viral genome; inverse PCR and comparison with known subtelomere sequences; long-read sequencing; optical genome mapping (Daibata *et al*., 1999; Huang *et al*., 2014; Luppi *et al*, 1998; Nacheva *et al*, 2008) (Aswad *et al*., 2021; Greninger *et al*., 2018; Kawamura *et al*., 2017; Liu *et al*., 2020; Miura *et al*., 2018; Tweedy *et al*., 2016; Wight *et al*, 2020; Zhang *et al*., 2017).

Detection of viral gene expression in some iciHHV-6A/6B carriers has raised the possibility of a deleterious impact on the individual’s health over their lifetime (Huang *et al*., 2014; Peddu *et al*, 2019; Strenger *et al*, 2014). Similarly, the presence of a large viral genome within in a telomere has been shown to cause localised instability that may influence telomere DNA damage signalling and function (Huang *et al*., 2014; Wood & Royle, 2017; Zhang *et al*, 2016). Recent studies have also indicated an association between iciHHV-6A/6B and angina pectoris (Gravel *et al*, 2015); unexplained infertility (Miura *et al*, 2020); an increased risk of aGVHD following haematopoietic stem cell transplantation when either the donor or recipient has iciHHV-6A/6B (Hill *et al*, 2017; Weschke *et al*, 2020); complications following solid organ transplantation when there was iciHHV-6A/6B donor/recipient mismatch (Bonnafous *et al*, 2018; Bonnafous *et al*, 2020; Petit *et al*, 2020); and possibly an increased risk of pre-eclampsia (Gaccioli *et al*, 2020).

There is increasing evidence that iciHHV-6A/6B genomes can reactivate fully (Endo *et al*, 2014) and then be transmitted as acqHHV-6A/6B (Gravel *et al*, 2013; Hall *et al*, 2010), but the chain of events leading to reactivation is unknown. Telomeres are terminated by a single-stranded 3′ overhang that can strand-invade into the upstream double-stranded telomeric DNA to form a telomere loop (t-loop) (Doksani *et al*, 2013; Griffith *et al*, 1999; Wang *et al*, 2004). This secondary structure is stabilised by TRF2, a component of the Shlerterin complex that binds to double-stranded telomere repeats, and plays a protective role by preventing the ends of linear chromosomes being detected as double-strand breaks and inappropriately repaired (de Lange, 2018; Schmutz *et al*, 2017; Stansel *et al*, 2001; Van Ly *et al*, 2018). However, t-loops can be excised as telomere repeat-containing t-circles (Pickett *et al*, 2009; Pickett *et al*, 2011; Sarek *et al*, 2015; Tomaska *et al*, 2019) by the SLX1-4 complex, which is a structure-specific endonuclease (Fekairi *et al*, 2009; Vannier *et al*, 2012; Wan *et al*, 2013), and unregulated excision of t-loops results in critically short telomeres (Deng *et al*, 2013; Pickett *et al*., 2011; Rivera *et al*, 2017). We and others have proposed that release of the viral genome is a prerequisite for reactivation of iciHHV-6A/6B and that this release is driven by normal t-loop processing (Huang *et al*., 2014; Prusty *et al*, 2013; Wood & Royle, 2017).

Here, we have investigated iciHHV-6A/6B genomes (with a particular focus on iciHHV-6B), their associated telomeres and their relationships to acqHHV-6B genomes in families and communities. We exploited the <u>p</u>roximal hyper<u>v</u>ariable region of <u>T1</u> (pvT1) in the HHV-6B genome to distinguish between acqHHV-6B strains and to predict relationships between iciHHV-6B and acqHHV- 6B in families. For the first time, we have detected acqHHV-6B telomeric integration in saliva DNA from healthy adults. The frequency of partial or complete iciHHV-6B excision varied between germline and somatic cells, with notably high frequencies in two pluripotent cell lines, and we present evidence of HHV-6B transmission from an iciHHV-6B carrier mother to her non-iciHHV-6B son. In summary, our study demonstrates that HHV-6B can readily transition between iciHHV-6B and acqHHV-6B forms, and vice versa.

## Results

### DRR-pvT1 tracking of HHV-6B transmission

To explore the relationship between iciHHV-6A/6B and acqHHV-6A/6B, we identified a genomic marker (pvT1) capable of distinguishing viral strains by nested PCR and sequencing. To investigate DRR-pvT1 specifically, the first round of amplification was achieved using a primer anchored at the right end of U near DRR and a primer in DR on the other side of T1 (Figure 1A). The second round of amplification and subsequent sequencing involved a primer (TJ1F) anchored in a short, conserved sequence approximately 400 bases into the T1 telomere-like repeat array and a second primer in DR. A total of 102 DNA samples were analysed: iciHHV-6B from 39 individuals (38.2%) and acqHHV-6B from 63 (61.8%) individuals (Supplementary Table 1). The analysis showed that DRR-pvT1 can be divided into three regions (left, middle and right), with most of the variation being due to differences in the number and distribution of telomere (TTAGGG) and degenerate telomere-like repeats (predominantly CTAGGG and CTATGG) in the left and middle regions. Maps of the repeat pattern are shown for a subset of samples in Figure 1B, and the complete dataset is provided in Figure 1 – figure supplement 1.

**Figure 1.**
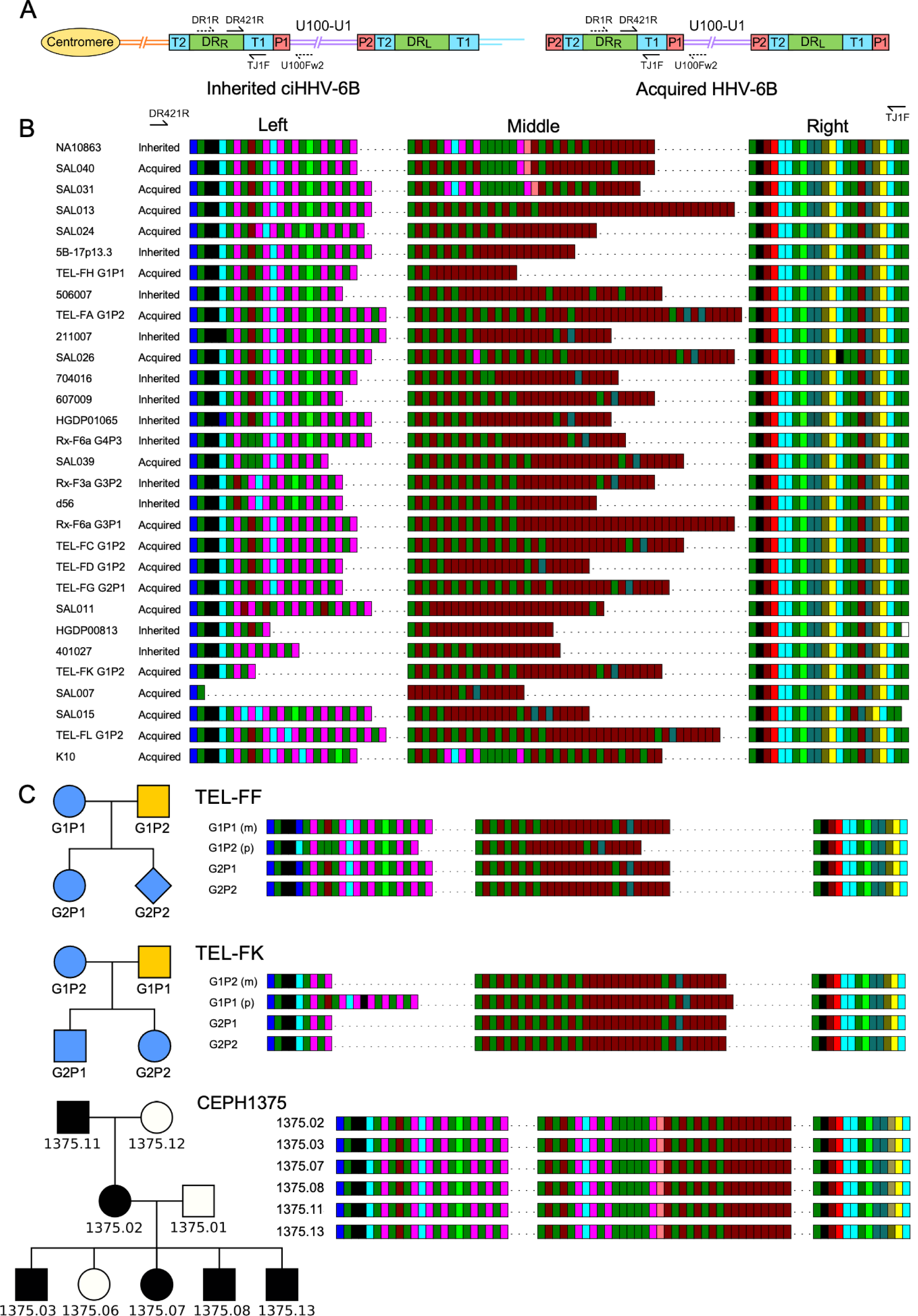
Characterisation of the highly variable DRR-pvT1 in iciHHV-6B and acqHHV-6B genomes. (A) Diagram showing the locations of PCR primers used to amplify the DRR-T1 region specifically (U100Fw2 and DR1R) and the nested primers used to reamplify and sequence pvT1 (DR421R and TJ1F) in the iciHHV-6B genome (left) and in the acqHHV-6B genome (right). (B) DRR-pvT1 repeat patterns are shown to demonstrate diversity among a subset of iciHHV-6B (inherited) and acqHHV- 6B (acquired) genomes from various individuals. Telomere-like and degenerate repeats present in the HHV-6B pvT1 region are colour-coded: Dark green, TTAGGG; brown, CTAGGG; cyan, TTAGTG; yellow, TTACTG; dark yellow, ATAGAC; teal, CTAAGG; pink, CTATGG; lime green, TTATGG; blue, GTAGTG; peach, TTAGAG; red, GTCTGG. Black squares represent other less common degenerate repeats, white squares show where the sequence could not be determined accurately. Dashes between repeats were added to maximize alignment and to allow comparison between the left, middle (highly variable) and right (highly conserved) sections of DRR-pvT1 in different samples. (C) DRR-pvT1 repeat patterns from acqHHV-6B in two families. None of the family members were iciHHV-6 carriers. In both families the HHV-6B DRR-pvT1 region in the two children was the same as one parent, indicating the children carried the same strain of the virus, whereas the second parent carried a different strain. Blue shapes indicate that the child had acqHHV-6B with the same pvT1 repeat map as the mother. Analysis of DRR-pvT1 in the CEPH1375 family showed stable inheritance of the same pvT1 repeat pattern across three generations. iciHHV-6B carriers in the CEPH1375 family are shown as filled black symbols

Among the 102 repeat patterns, 93 were different from each other, and 90 of these were detected in only a single donor or family. Amongst the 12 repeat patterns shared between individuals, eight were identical to each other, and seven of these were from closely related viral genome sequences in iciHHV-6B carriers with a shared 9q integration. The eighth was found in saliva DNA from an individual (SAL030) with acqHHV-6B (Figure 1 – figure supplement 1). Among the remaining four repeat patterns, two were in iciHHV-6B samples (NWA008 and DER512) that share the same 17p integration site, and two were in acqHHV-6B samples (SAL023 and TEL-FA G1P1).

Remarkably, the pvT1 repeat pattern was different in almost every unrelated individual with acqHHV-6B (61/63, 96.8%) and also in the majority of iciHHV-6B individuals (30/39, 77.0%; 29/31, 93.5%, when the entire 9q iciHHV-6B group was excluded).

Although DRR-pvT1 is highly variable among acqHHV-6B genomes in unrelated individuals, this is not the case within families (Figure 1C, Figure 1 – figure supplement 2). Saliva DNA was analysed from 36 individuals in eight families (without iciHHV-6A/6B) and DRR-pvT1 was successfully amplified and sequenced from 34 of them (94%). Eighteen of the 20 children (90%) in these families had the same repeat pattern as one of their parents, presumably as a result of acqHHV-6B transmission directly from a parent (maternal transmission in six of the eight families and paternal transmission in one) or indirectly via a sibling (Figure 1 – figure supplement 2). In contrast, the analysis revealed that one child in each of two families must have contracted acqHHV-6B from outside their immediate family (TEL-FC G2P2 and TEL-FM G2P1, Figure 1 – figure supplement 2). As a control, analysis of DRR-pvT1 repeat patterns in iciHHV-6B carriers in the CEPH1375 family demonstrated the stability of the repeat patterns across three generations (Figure 1C). All six individuals, shown previously to be iciHHV-6B carriers (Huang *et al*., 2014), shared an identical pattern indicating that DRR-pvT1 does not necessarily acquire mutations from one generation to the next. In summary, DRR-pvT1 analysis was used successfully to distinguish HHV-6B strains and to monitor transmission of iciHHV-6B and acqHHV-6B between family members.

### Phylogeny of iciHHV-6A/6B in relation to integration events

Progress has been made recently in sequencing iciHHV-6A/6B and acqHHV-6A/6B genomes by generating overlapping PCR amplicons (Zhang *et al*., 2017) or an oligonucleotide hybridisation enrichment process (Aswad *et al*., 2021; Greninger *et al*., 2018; Telford *et al*, 2018). However, the evolutionary histories of iciHHV-6A/6B genomes and their relationships with acqHHV-6A/6B genomes remain unclear. Moreover, it is not known how many founder germline integration events have occurred. To expand the data required to answer such questions, we screened various cohorts and families to identify additional iciHHV-6A/6B-positive samples (Supplementary Table 1), and then selected several key samples for viral genome sequencing using the overlapping PCR amplicon approach (Huang *et al*., 2014; Zhang *et al*., 2017). These samples included five iciHHV-6A and four iciHHV-6B samples from the BIOSTAT-CHF cohort, four other iciHHV-6A samples, one iciHHV-6A cell line (HGDP00628 (Huang *et al*., 2014)), and a pluripotent cell line (d37) from an iciHHV-6B carrier. In total, ten iciHHV-6A and five iciHHV-6B genomes were sequenced.

Distance-based phylogenetic networks of iciHHV-6A/6B and acqHHV-6A/6B genomes were generated, including the 15 newly sequenced iciHHV-6A/6B genomes and iciHHV-6A/6B genomes representing previously identified clades and integration sites (Figure 2, Supplementary Table 1). These networks reinforce previous findings (Aswad *et al*., 2021; Greninger *et al*., 2018; Tweedy *et al*., 2016; Zhang *et al*., 2017) showing that the majority of iciHHV-6A genomes in European and North American carriers (including the ten newly sequenced) originate from three integration events at 17p, 18q and 19q, with the time to the most recent common ancestor estimated at 23 to 105 thousand years ago (Figure 2, Supplementary Table 2). The networks also confirm that iciHHV-6B carriers with European and North America ancestry originate from a larger number of integration events that occurred more recently (21 to 25 thousand years ago; Supplementary Table 2) (Aswad *et al*., 2021; Greninger *et al*., 2018; Zhang *et al*., 2017). Inclusion of the newly sequenced iciHHV-6B genomes added two new iciHHV-6B clades to the phylogenetic network, each defined by a distinct integration site (Figure 2). As a consequence of the complex evolution of subterminal sequences in the human genome (Riethman, 2008), the chromosomal locations in these iciHHV-6B clades will require further verification by an independent method, but they represent two more independent germline telomeric integration events.

**Figure 2.**
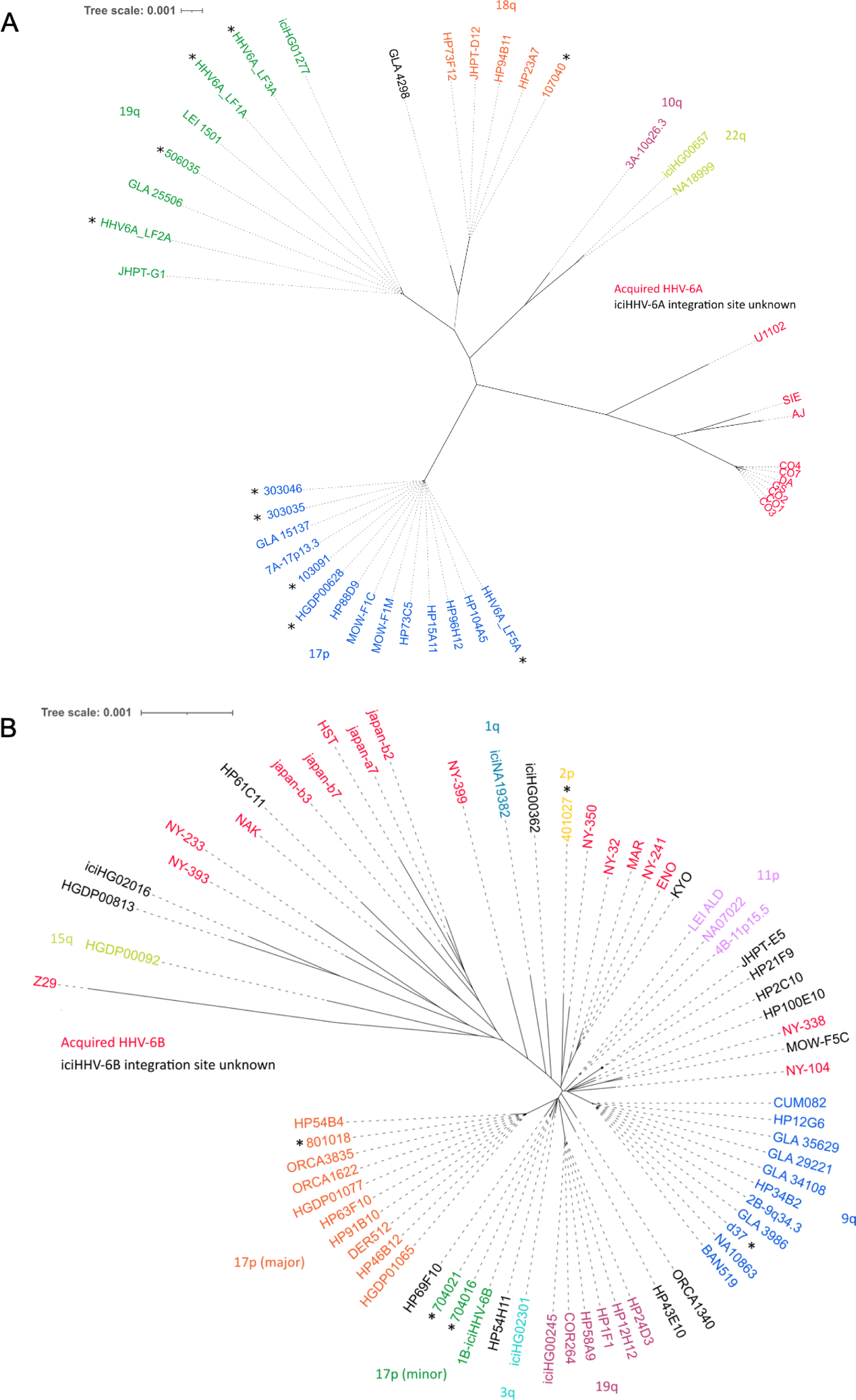
Distance-based phylogenetic networks for HHV-6A and HHV-6B. (A) A phylogenetic network of 41 HHV-6A viral genome sequences. High sequence homology to iciHHV-6A genomes for which the integration site had already been established allowed all but one (GLA_4298) iciHHV-6A genomes to be assigned to a telomeric integration site. The colours indicate an chromosomal integration as follows: Green, 19q; Orange, 18q; Purple, 10q; Yellow-green, 22q; Blue, 17p. The name in black identifies the iciHHV-6A strain without a predicted chromosomal location. Non-inherited, acqHHV-6A strains are shown in red. iciHHV-6A genomes sequenced as part of this study are identified by a black asterisk. (B) A phylogenetic tree of 68 HHV-6B genome sequences (51 iciHHV-6B and 17 acqHHV-6B). iciHHV-6B genomes sequenced as part of this study are identified by a black asterisk. High sequence homology to iciHHV-6B genomes with a known integration site (already established by FISH) allowed the majority of iciHHV-6B genomes to be assigned a telomeric chromosomal location. From the newly sequenced iciHHV-6B genomes two new clades were identified and provisionally labelled as: 17p (minor) and 2p. The 17p (minor) clade includes the 1B- iciHHV-6B genome, previously a singleton in HHV-6B networks, and the iciHHV-6B genomes in 704021 and 704016. The subtelomere-iciHHV-6B junction sequences, amplified with the 17p311 and DR8F(A/B) primers, are similar for all three samples and distinct form the 17p (major) clade. The other new clade is characterised by the iciHHV-6B genome (401027). This group is provisionally labelled 2p because the subtelomere-iciHHV-6B junction was amplified by the 2p2 and DR8F(A/B) primers in 401027, and two other samples (410005 and 308006, see Figure 3C).

As an aid to analysing the increasing number of HHV-6A/6B genome sequences, we developed HHV-6 Explorer (https://www.hhv6explorer.org/), which is an online interface for monitoring clade-specific variation in DNA and predicted protein sequences. An analysis of sequence variation in relation to integration site using this tool produced new insights, for example by demonstrating that two potentially inactivating mutations, U14 in an iciHHV-6B clade and in U79 in an iciHHV-6A clade, must have arisen after integration (Figure 2 – figure supplement 1).

### Predicting chromosomal iciHHV-6B integration sites from DRR-pvT1 repeat patterns

To explore whether DRR-pvT1 repeat patterns reflect the phylogenetic relationships between iciHHV-6B genomes, 19 for which DNA was available, were subjected to DRR-pvT1 analysis. The results show that iciHHV-6B genomes with a high degree of overall sequence similarity also shared similar DRR-pvT1 repeat patterns. For example, individuals with 9q iciHHV-6B have a characteristic CTATGG- TTAGTG-CTATGG motif in the middle region of DRR-pvT1, as well as a TTAGAG repeat that is found infrequently outside this group. Also, viral sequences assigned to the 17p major iciHHV-6B group by genome sequence identity and shared 17p subtelomere-iciHHV-6B junction sequences (HGDP01065, HGDP01077, DER512, 801018 and ORCA1622) (Zhang *et al*., 2017) had similar DRR-pvT1 repeat patterns with two characteristic GTAGTG repeats at the left end (Figure 3A).

**Figure 3.**
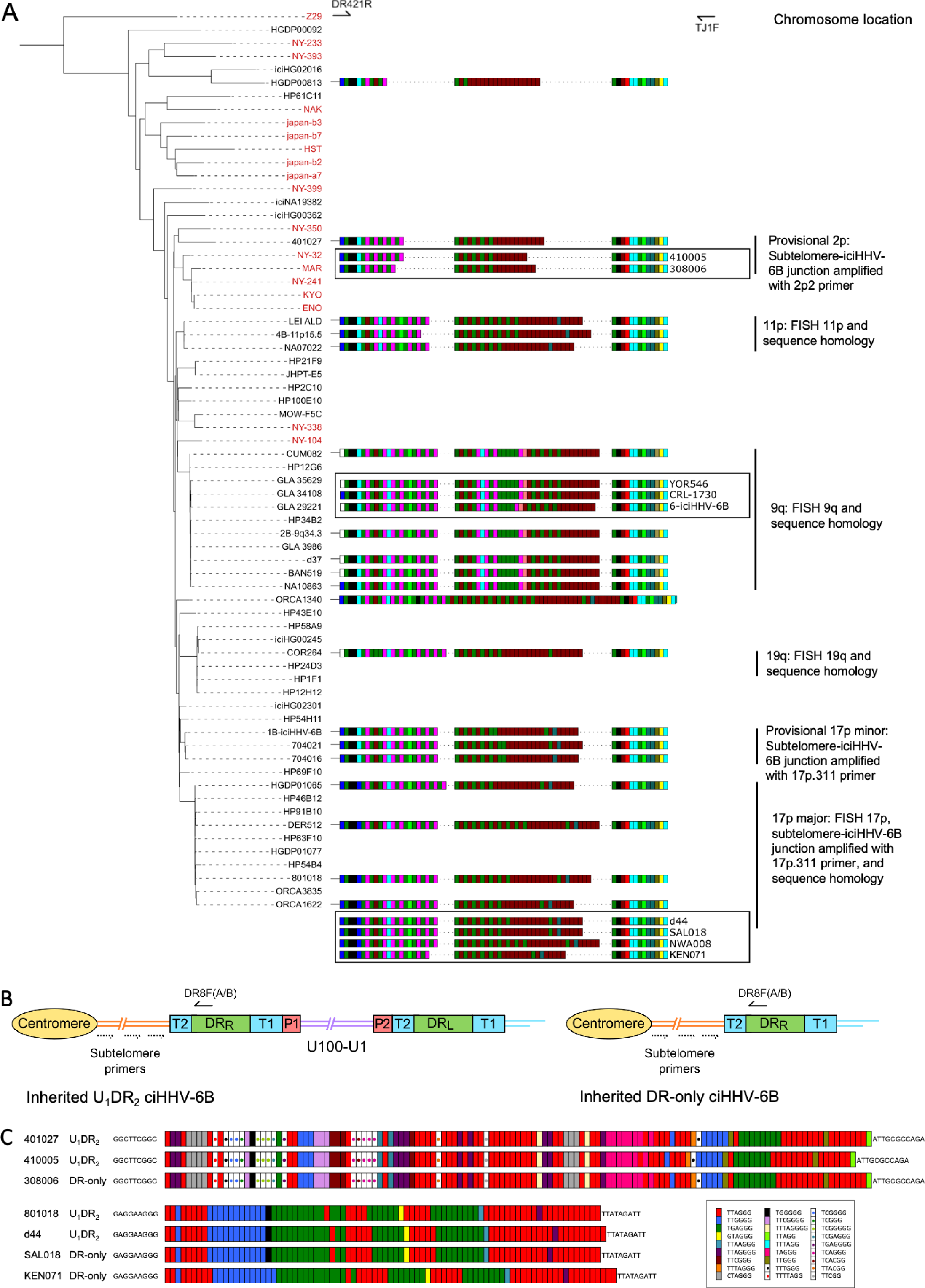
pvT1 repeat patterns are similar between strains of iciHHV-6B known to have the same ancestral integration site and those predicted to share that integration site. (A) Distance-based phylogenetic tree of selected iciHHV-6B (black names) and acquired HHV-6B (red names) genomes. Where DNA was available, DRR-pvT1 repeat patterns from 19 iciHHV-6B samples in the phylogenetic tree are shown. The repeats in the pvT1 region are colour-coded: Dark green, TTAGGG; brown, CTAGGG; cyan, TTAGTG; yellow, TTACTG; dark yellow, ATAGAC; teal, CTAAGG; pink, CTATGG; lime green, TTATGG; blue, GTAGTG; peach, TTAGAG; red, GTCTGG. Black vertical bars on the right indicate groups of iciHHV-6Bs for which the integration site has been determined by FISH, phylogeny, subtelomere-iciHHV-6B junction analysis or a combination of these. DRR-pvT1 repeat patterns within black boxes identify iciHHV-6B samples for which the integration site has been predicted based on DRR-pvT1 similarity. For some of these samples the integration site has been validated by subtelomere-iciHHV-6B junction analysis (410005, 308006, d44, SAL018, NWA008 and KEN071). (B) Line diagram showing the position of DR8F(A/B) and potential subtelomere primers used to amplify and sequence subtelomere-iciHHV-6B junctions. The subtelomere-iciHHV-6B integration site can be amplified whether the integrated viral genome is full length or iciHHV-6B-DR only. (C) Diagram showing of the pattern of canonical (TTAGGG) and degenerate telomere-like repeat maps across the subtelomere-HHV-6B junction from two different chromosome ends. The subtelomere-HHV-6B repeat maps from the 401027, 410005 and 308006 were generated following amplification with primers 2p2 and DR8F(A/B). They are very similar demonstrating common ancestry (top three maps). The subtelomere-HHV-6B repeat maps from the 801018, d44, SAL018 and KEN071 generated by amplification with 17p311 and DR8F(A/B) (bottom four maps) are very similar to the group of iciHHV-6B genomes in the 17p (major) clade some of which have been shown to be integrated in 17p. The 308006, SAL018 and KEN071 samples contain iciHHV-6B DR-only.

From these observations, we hypothesised that DRR-pvT1 repeat patterns may be used to predict both the integration site for newly identified iciHHV-6B carriers and the identity of closely related viral strains, without the need to sequence viral genomes. Based on similarity between DRR- pvT1 repeat patterns, the integration site was predicted for nine iciHHV-6B genomes for which genome sequences were not available (Figure 3A). For example, the repeat patterns in YOR546, CRL- 1730 and 6-iciHHV-6B were identical or almost identical to those in a large group of iciHHV-6B genomes with a 9q integration site (Nacheva *et al*., 2008; Shioda *et al*, 2018). PCR amplification of the subtelomere-iciHHV-6B junction was used to validate several of the predictions arising from these results. For example, DRR-pvT1 analysis placed 410005 and 308006 in the same phylogenetic clade as the 401027 iciHHV-6B genome (Figure 3, Figure 2), and common ancestry was confirmed by sequence similarity across the subtelomere-iciHHV-6B junction fragment (amplified by primer 2p2, Figure 3B-C, Figure 2) in all three samples. Similarly, the iciHHV-6B genomes in d44, LAT018, NWA008 and KEN071 share DRR-pvT1 repeat patterns and subtelomere-iciHHV-6B junction fragments similar to the large group of iciHHV-6B genomes integrated at 17p (17p major) (Figure 3B- C, (Zhang *et al*., 2017)). In summary, the integration sites of iciHHV-6B genomes that had not been sequenced were predicted based on a high degree of similarity between DRR-pvT1 repeat patterns, and six of these predictions were subsequently tested and validated by an independent method.

### Chromosomal integration of acqHHV-6B

Telomeric integration provides a means by which acqHHV-6A/6B may achieve latency.

Although *de novo* integration has been shown to occur in cell culture (Arbuckle *et al*., 2010; Arbuckle *et al*, 2013; Collin *et al*, 2020; Gilbert-Girard *et al*., 2020; Gravel *et al*, 2017; Wallaschek *et al*., 2016b), there is little evidence that it occurs contemporaneously *in vivo* (Moustafa *et al*, 2017). To investigate this, we collected saliva DNA samples from 52 healthy donors and used triplicate digital droplet PCR (ddPCR) assays to quantify HHV-6B genome copy number per cell (Figure 4A). Using this sensitive approach, low-level HHV-6B was detected in 44 samples (84.6%) with a mean value 0.00145 copies per cell (range 0.0000233–0.0125). Two kidney DNA samples, K1 and K10, were also positive for HHV-6B at 0.000525 and 0.0124 copies per cell, respectively (Figure 4A, D).

**Figure 4.**
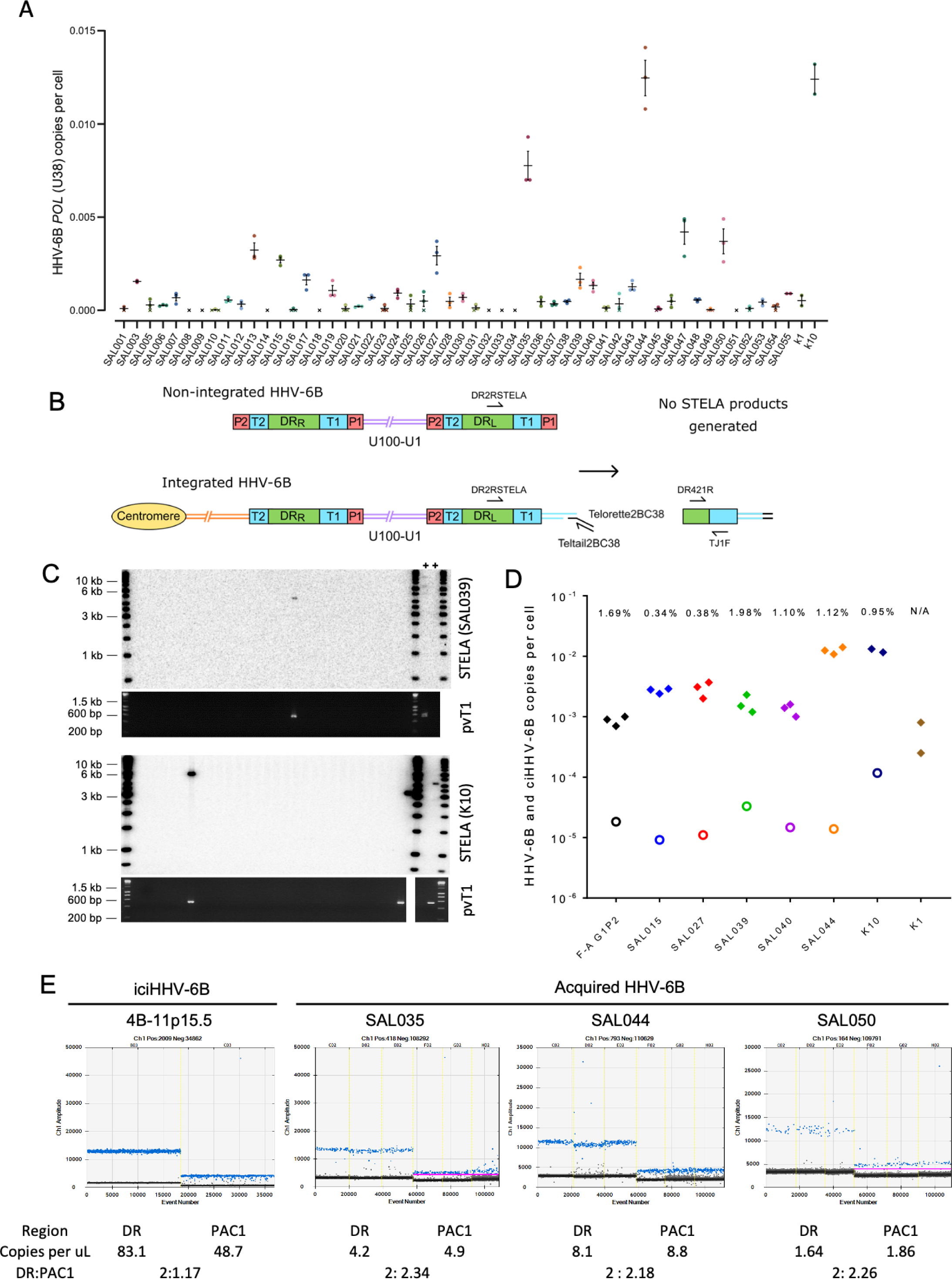
Evidence of telomere formation at the distal end of HHV-6B genomes in individuals with community-acquired HHV-6B. (A) Graph showing HHV-6B copy number per cell estimated from HHV-6B *POL* (U38) and RPP30 copy number by ddPCR. Triplicate results are shown for saliva DNA from 52 individuals and kidney samples from two individuals. Crosses show where zero copies per cell were estimated for a single replicate and black crosses indicate that zero copies per cell were estimated from all three replicates. The mean and standard error are shown. (B) STELA primers, DR2RSTELA and Telorette2BC38/TeltailBC38, that were used to amplify HHV-6B associated telomeres are shown on an integrated HHV-6B genome. DR421R and TJ1F primers were used to amplify DRL-pvT1 in secondary nested PCRs. In non-integrated HHV-6B STELA products cannot be generated as there is no HHV-6B-associated telomere. The copies of PAC1 (P1) and PAC2 (P2) are shown. (C) STELA blots from two DNA samples (SAL039 and K10) are shown. In a small number of reactions an amplified HHV-6B-associated telomere was detected with a radiolabeled (TTAGGG)n telomere-repeat probe. Diluted STELA PCR products were used as input for secondary, nested PCRs to amplify DRL-pvT1 as shown in the agarose gel photograph below. The 6-iciHHV-6B DNA was used as a positive control (+) for STELA and DRL-pvT1 secondary PCR. (D) Graph showing the total copy number of HHV-6B per cell measured by ddPCR (filled diamonds); the estimated number of copies of integrated HHV-6B per cell from STELA and pvT1 PCR (open circles). (E) Measuring the DR:PAC1 ratio by ddPCR. 1D EvaGreen ddPCR plots of one iciHHV-6B (4B-11p15.5) cell line and acqHHV-6B in three saliva samples (SAL035, SAL044 and SAL050). Triplicate reactions for a DR amplicon (DR6B-F and DR6B-R primers) and a PAC1 amplicon (PAC1F and PAC1R-33 primers) are shown. The absolute copy number of each amplicon per μl ddPCR reaction and the ratio between DR and PAC1 amplicons are shown below each plot.

Previously we used Single TElomere Length Analysis (STELA), a PCR-based method, to detect and measure the length of iciHHV-6A/6B-associated telomeres at DRL-T1 (Huang *et al*., 2014). To determine whether some copies of acqHHV-6B are integrated into telomeres, we aimed to detect these potentially rare events using STELA. The first step involved STELA of genomic DNA from saliva samples that had low but measurable levels of HHV-6B DNA. The STELA PCR products were then diluted and subjected to a second round of PCR with primers TJ1F and DR421R to amplify DRL-pvT1. Successful amplification in the second round was taken as evidence that an HHV-6B-associated telomere had been amplified in the primary STELA round (Figure 4B-C). Sanger sequencing was used to confirm DRL-pvT1 amplified from STELA products matched that amplified from genomic DNA from each individual. Various steps were taken to avoid false positive results. These include the use of newly designed STELA primers for these experiments (Figure 4B) and conducting STELA on genomic DNA from the HT1080 cell line (HHV-6A/6B negative) to check for potential non-specific amplification from another telomere or elsewhere in the human genome. No STELA products were generated from HT1080 among 900 STELA reactions, total 4.5μg DNA screened. This two-step STELA procedure, conducted on DNA from six saliva and two kidney samples identified a small number of amplicons from some samples with acqHHV-6B (Supplementary Table 3). The proportion of HHV-6B genomes from which a telomere was amplified was estimated using the HHV-6B copy number per cell (determined by ddPCR) and by estimating the number of cell equivalents of DNA used per STELA reaction, assuming 6.6 pg DNA per cell (Figure 4D, Supplementary Table 3). For the eight samples analysed, an average of 0.95% (range 0–1.98%) of HHV-6B genomes resulted in pvT1 amplification, indicative of telomeric integration. The low copy number of HHV-6B in these samples, combined with reduced PCR amplification efficiency of longer molecules (telomeres in this case), has a stochastic effect on the potential to detect an HHV-6B-associated telomere in each STELA reaction therefore, the integration frequencies should be interpreted cautiously.

We then sought to measure the proportion of acqHHV-6B genomes that are *not* integrated in saliva samples, using ddPCR to quantify the copy numbers of DR and PAC1. PAC1 and PAC2 are genome packaging signals located at the end of T1 and T2, respectively (Figure 4B). The linear, unintegrated genome has two copies of DR and two copies of PAC1 and PAC2, whereas the integrated genome has two copies of DR but retains only a single copy of PAC1 and PAC2 (Figure 4B) (Arbuckle *et al*., 2010) (Huang *et al*., 2014). As expected, the ratio of DR:PAC1 in an iciHHV-6B sample (4B-11p15.5) was approximately 2:1, whereas the ratio in acqHHV6-B samples in saliva samples was approximately 2:2 (Figure 4E). This shows that most acqHHV-6B genomes in saliva samples are not integrated, consistent with the STELA data that revealed only low level telomeric integration of acqHHV-6B genomes in saliva DNA samples.

### Partial excision of iciHHV-6B genomes and novel telomere formation at DRL-T2

We showed previously that iciHHV-6A/6B genomes in lymphoblastoid cell lines (LCLs) can be truncated at DRL-T2, resulting in loss of the terminal DRL and formation of a novel short telomere at the breakpoint (Huang *et al*., 2014; Wood & Royle, 2017). We proposed that truncation is achieved through a t-loop-driven mechanism in which the 3′ G-rich overhang at the end of the telomere strand invades DRL-T2, facilitating excision of the terminal DRL as a DR-containing t-circle and leaving a novel telomere at the excision point. Here, we measured the frequency of iciHHV-6B truncations at DRL-T2 in seven LCLs, nine blood DNAs (from white blood cells), two pluripotent cell lines, three sperm DNA samples and saliva DNA (Figure 5A-C, Supplementary Table 4). The average frequency of DRL-T2 truncations with novel telomere formation was 1 in 120 cells, but there were significant differences between DNAs from different cell types. The number of truncations per cell was lowest in blood and saliva DNA, with an average of 1 in 300 (0.0033 per cell) and 1 in 320 (0.0031), respectively, and approximately double this in sperm at 1 in 160 cells (0.00627 per cell). An astonishingly high number of DRL-T2 truncations were observed in LCLs (0.015 per cell) and pluripotent cells (0.014 per cell), equating to approximately one in every 70 cells. Furthermore, released circular DR molecules were detected using an inverse PCR approach (Figure 5 – figure supplement 1).

**Figure 5.**
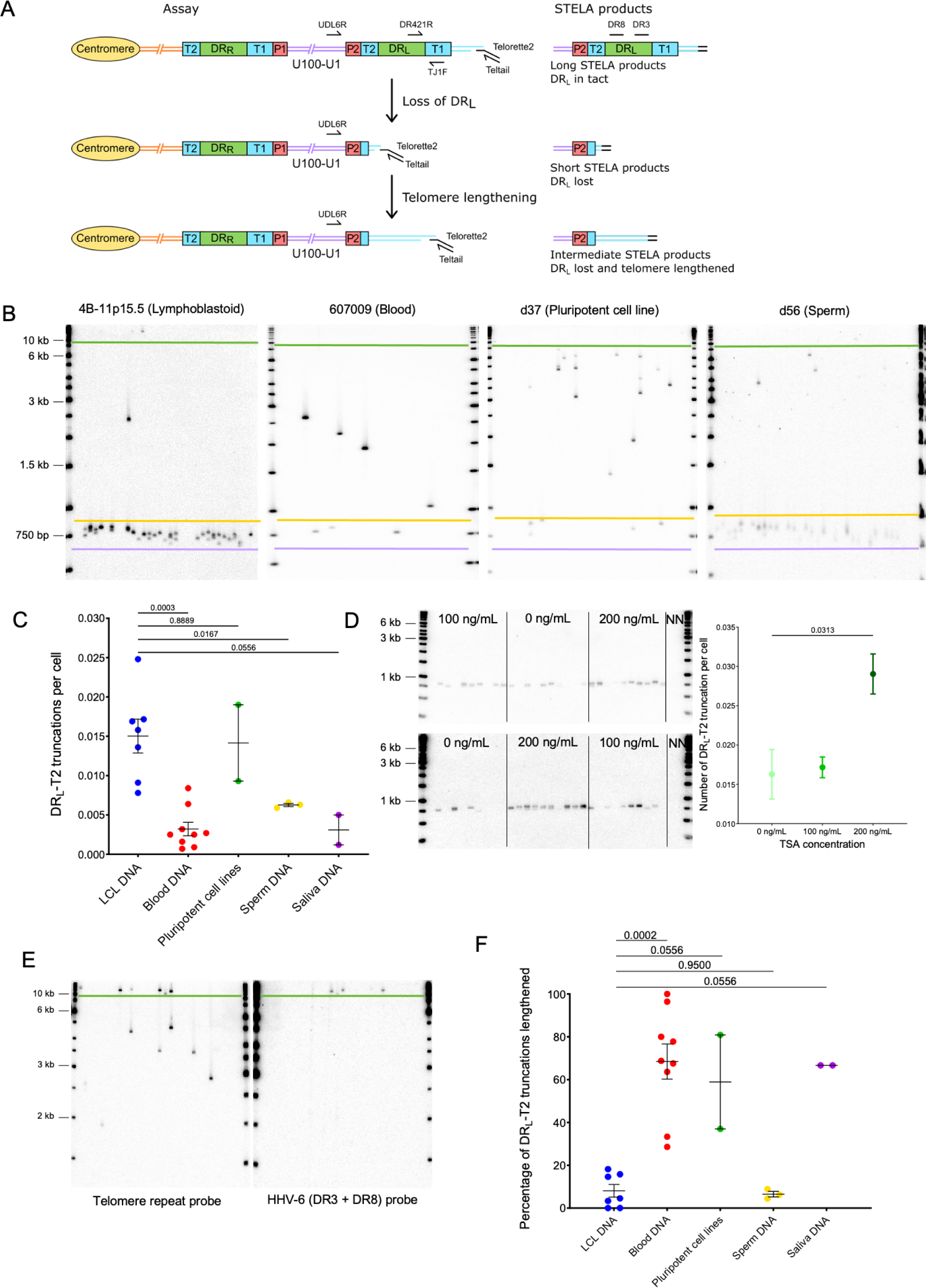
Quantifying iciHHV-6B truncations at DRL-T2 and assessment of the newly formed telomere length. (A) Diagram showing the loss of DRL from iciHHV-6B, formation of a novel telomere at DRL-T2 and the potential lengthening of the newly formed telomere by a telomere maintenance mechanism. STELA primers (Teltail and Telorette2) and the primer UDL6R, in the unique region of the iciHHV-6B genome, were used to amplify HHV-6B-associated telomeres. Long STELA products (>8.6kb) are generated from intact copies of the iciHHV-6B genome and include amplification through DRL to the end of the telomere. Short STELA products (0.6 - 0.9 kb) are generated when DRL has been lost and a novel telomere has formed at DRL-T2. Intermediate sized STELA products are generated when the novel telomere formed at DRL-T2 has been lengthened. All three types of STELA amplicons contain TTAGGG telomere repeats and hybridise to a telomere repeat probe. The relative positions of the DR3 and DR8 probes used to detect DR sequences are shown. (B) Representative HHV-6B STELA blots generated from different cell-types from iciHHV-6B carriers or derived cell lines. These include a lymphoblastoid cell line (LCL) from individual 4B-11p15.5; white blood cells from individual 607009; a pluripotent cell line d37; sperm DNA d56. Bands between the mauve and yellow lines are short STELA products. The bands detected between the yellow and green lines are intermediate length STELA products, representative of newly formed telomeres at DRL-T2 that have been lengthened. Long STELA amplicons, above the green line, are products generated from full- length copies of iciHHV-6B and include DRL. (C) Graph showing the estimated number of DRL-T2 truncations per cell, including those that have been lengthened, in DNA from different iciHHV-6B individuals and cell types. Means with standard error and the p-values from a Mann-Whitney ranked sum test to compare truncation frequencies (LCL vs other cell-types) are shown. (D) Representative UDL6R-STELA blots from the 4B-11p15.5 cell line treated with 0, 100 and 200 ng/mL of trichostatin-A and a graph showing the frequency of truncations per cell (mean and standard error). Data were derived from UDL6R-STELA conducted twice (technical replicates) using DNA extracted from treated and untreated cells grown in triplicate (biological replicates). p-values were from a Wilcoxon test. (E) DRL-T2 truncation assay on blood DNA from the 401027 iciHHV-6B carrier demonstrating that the long STELA products (above the green line) contain DR sequences that hybridise to the telomere repeat probe (left panel) and the combined DR3/DR8 probe (right panel). Whereas the intermediate length amplicons (below the green line) only hybridise to the telomere repeat probe (left panel) demonstrating that each amplicon represents newly formed telomere at DRL-T2 that has been lengthened. (F) The graph shows the percentage of newly formed telomeres at DRL-T2 that were lengthened in samples from various unrelated iciHHV-6B carriers and cell types. Means with standard error and the p-values from a Mann-Whitney ranked sum test to compare truncation frequencies (LCL vs other cell-types) are shown.

The histone deacetylase inhibitor, trichostatin-A (TSA) has been shown to promote iciHHV- 6A reactivation in cell lines and cultured T cells from iciHHV-6A/6B individuals (Arbuckle *et al*., 2010; Arbuckle *et al*., 2013), and also to increase the abundance of t-circles (Zhang *et al*, 2019). To explore whether the frequency of truncations with short telomere formation at DRL-T2 could be influenced by the chromatin state of the iciHHV-6B genome, the 4B-11p15.5 lymphoblastoid cell line was treated with TSA. The cells were grown in medium supplemented with various concentrations of TSA for approximately 5 days, and the DRL-T2 truncation assay (DRL-T2-STELA) was conducted on DNA extracted from TSA-treated and control cells (Figure 5D). The frequency of truncation events at DRL- T2 increased significantly from 0.0162 to 0.0290 per cell at the highest TSA concentration. This suggests that iciHHV-6B chromatin conformation influences the chance of t-loop formation at the telomere-like repeats within the viral genome and subsequent excision events.

Loss of the terminal DRL via t-loop formation and excision at DRL-T2 was expected to generate a short novel telomere detected as a STELA fragment consisting of a flanking region of DRL of approximately 600 bp plus a telomere repeat array limited to the length of the DRL-T2 (Figure 5A). Some of the DRL-T2-STELA amplicons were larger than the expected 0.7-0.9 kb (Figure 5B). We showed that these amplicons lack DR3 or DR8 sequences (Figure 5E), thus indicating they are not molecules truncated at different sites within DRL. We hypothesized that the intermediate length amplicons (longer than 0.7-0.9kb, Figure 5B) could have arisen by telomerase-mediated lengthening of newly formed telomeres at the DRL-T2 truncation site. To address this, six of the lengthened amplicons from three DNA samples (4B-11p15.5, d56 and NWA008) were sequenced. All comprised the expected flanking sequence followed by a uniform array of (TTAGGG) repeats exceeding the known length of DRL-T2 for the sample, thus showing that some of the novel telomeres formed at DRL-T2 had been lengthened (Figure 5 – figure supplement 1). Telomerase activity was detected at a low level in the 4B-11p15.5 and other iciHHV-6B LCLs (Figure 5 – figure supplement 1), consistent with a telomerase-mediated lengthening of newly formed telomeres.

The percentage of DRL-T2 telomeres that had undergone lengthening varied between the iciHHV-6B samples and cell types (Figure 5F). The proportion of lengthened telomeres was higher in the pluripotent cell lines, blood DNA and saliva samples, and lower in the LCLs and sperm samples. This may reflect the level of telomerase activity in the cells or their progenitors, but it may also indicate variable responses to extremely short telomeres.

### Partial excision of iciHHV-6B genomes and novel telomere formation at DRR-T1

Processes driven by t-loop formation can also facilitate the release of U in its entirety with a single DR from iciHHV-6B (reconstituted with both PAC1 and PAC2) as a circular (U-DR) molecule (Arbuckle *et al*., 2013; Borenstein & Frenkel, 2009; Huang *et al*., 2014; Wood & Royle, 2017). The reciprocal product of U-DR excision is expected to be a truncation of the iciHHV-6B genome at DRR- T1 with new telomere formation. To explore this, we exploited the variable nature of pvT1.

Differences between DRR-pvT1 and DRL-pvT1 within a single iciHHV-6B genome were found in many of the iciHHV-6B samples analysed (24/35, 68.6%) (Figure 6A, Figure 6 – figure supplement 1). These differences were usually a consequence of loss or gain of a small number of repeats, although in some cases they involved single base substitutions that converted one repeat type to another. These differences made it possible to measure the frequency of truncations and new telomere formation at DRR-T1 via a two-step process.

**Figure 6.**
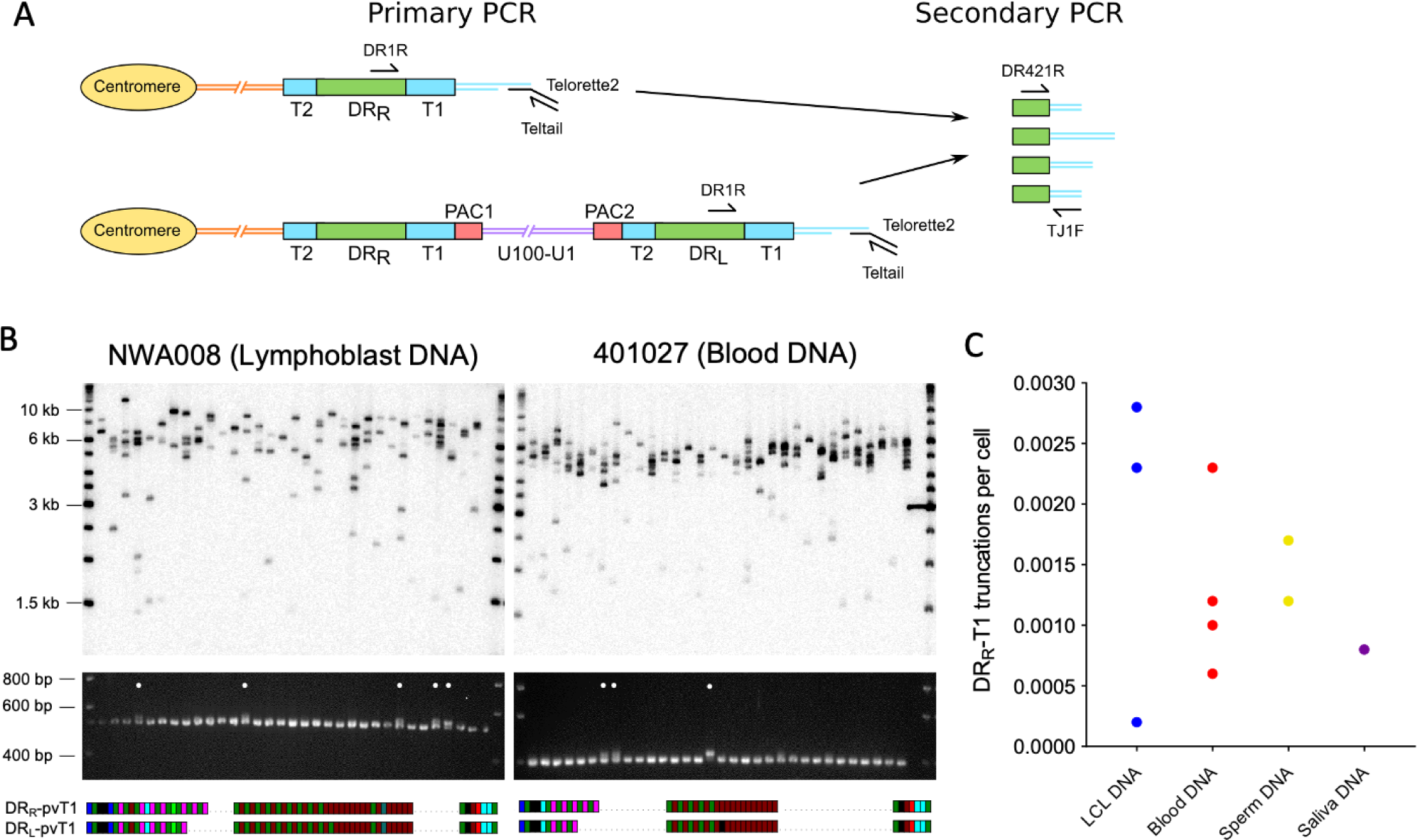
Measuring the frequency of iciHHV-6B truncations at DRR-T1 associated with novel telomere formation. (A) Schematic of the two-step assay used to differentiate between iciHHV-6B associated telomeres at DRR-T1 and DRL-T1. Telomeres at DRR-T1 and DRL-T1 were PCR amplified using the STELA primers, DR1R with Telorette2/Teltail. Subsequently, secondary nested PCRs were used to amplify pvT1 (primers DR421 and TJ1F) followed by size separation to distinguish pvT1 from DRR-T1 or DRL-T1. (B) Detection of telomeres at DRR-T1 or DRL -T1 in two iciHHV-6B samples (NWA008 and 401027) with known length differences between pvT1 in the DRs. Top images show telomeres at DRL and DRR in the iciHHV-6B samples amplified by STELA and detected by Southern blot hybridization to a radiolabeled (TTAGGG)n probe. Lower images show ethidium bromide-stained agarose gels of size separated pvT1 sequences amplified from the corresponding STELA reaction in the panel above. White dots identify the PCR reactions that contain pvT1 from DRR-T1. Below each set of panels are the pvT1 repeat maps from DRR-T1 or DRL-T1 for the corresponding sample, highlighting the differences. (C) Graph showing the frequency of truncations and new telomere formation at DRR-T1 in iciHHV-6B samples from unrelated individuals and cell types. The number of truncations per cell was calculated using the number of reactions from which DRR-pvT1 was amplified divided by the cell equivalents estimated from the total quantity of DNA screened (6.6 pg or 3.3 pg DNA per cell was assumed for diploid and haploid cells respectively).

First, the iciHHV-6B-associated telomeres were amplified using STELA with the DR1R primer that can generate products from a telomere at DRL-T1 (from normal, full-length iciHHV-6B) or DRR-T1 (following U-DR excision). Second, to distinguish between these STELA products, the PCR amplicons were subjected to nested PCR to amplify DRL-pvT1 and DRR-pvT1 (Figure 6A). The sizes of the pvT1 amplicons indicated whether the STELA products in the primary PCR had been derived from telomeres at DRL or DRR (Figure 6B), and selected sequencing was used to confirm. Truncations at DRR-T1 were detected in all ten iciHHV-6B samples analysed, and the frequency of newly formed telomeres at DRR-T1 was estimated to be 0.0002–0.0029 per cell (Figure 6C, Supplementary Table 5).

### Generation of DR-only iciHHV-6B by partial excision of full-length iciHHV6-B genomes in the germline

The frequency of truncations and new telomere formation at DRR-T1 in sperm DNA from iciHHV-6B carriers raised the strong possibility that such events could explain how individuals may inherit integrations consisting only of DR (DR-only iciHHV-6A/6B) (Huang *et al*., 2014; Liu *et al*., 2020; Ohye *et al*, 2014). To date, we have identified three individuals (308006 and SAL018 in the present study, and KEN071 (Huang *et al*., 2014)) who carry iciHHV-6B DR-only. Comparison of pvT1 repeat patterns in these samples with DRR-pvT1/DRL-pvT1 repeat patterns in full length iciHHV-6B suggested shared ancestry between these individuals with DR-only iciHHV-6B and those with full length iciHHV-6B (Figure 3A). To investigate this, attempts were made to amplify and sequence the subtelomere-iciHHV-6B junction, based on previously described junction fragments (Tweedy *et al*., 2016; Zhang *et al*., 2017) and using primers that anneal in the subtelomeric regions of various chromosomes (Figure 3B). Using this approach, we found that the DR-only iciHHV-6B in SAL018 and KEN071 share the common 17p (major) subtelomere-iciHHV-6B junction that is also found in 801018, d44 and others (Figure 3C). Similarly, the pattern of telomere and degenerate repeats across a subtelomere-iciHHV-6B junction (amplified by subtelomere primer 2p2) in the DR-only iciHHV-6B sample 308006, closely matched those in full length iciHHV-6B individuals 401027 and 410005 (Figure 3C). The existence of shared integration sites for the full-length and DR-only iciHHV- 6B carriers at two different chromosome ends clearly establishes that the DR-only status has arisen independently, on at least two occasions, by loss of U and one copy of DR in the germline of an ancestor with a full-length iciHHV-6B genome.

### Evidence of viral transmission from iciHHV-6B carrier mother to non-iciHHV-6B son

It has been shown that iciHHV-6A can reactivate in an immune-compromised setting (Endo *et al*., 2014) and there is evidence of transplacental transmission of reactivated iciHHV-6A/6B (Gravel *et al*., 2013). Despite these examples, the high prevalence of HHV-6B in populations has made it difficult to determine how often iciHHV-6B reactivation occurs. The evidence for frequent partial or complete release of iciHHV-6B genomes in somatic cells and the germline presented above suggests that opportunities for reactivation may be more common than currently appreciated. To explore this, we used DRR-pvT1 analysis to investigate the relationship between strains of iciHHV-6B and low-level acqHHV-6B within families. In one family (Rx-F6a), the mother (G3P1) had a low acqHHV-6B load in saliva (0.00035 copies per cell), but both her children were iciHHV-6B carriers.

The DRR-pvT1 repeat patterns in the children were identical and presumed to have been inherited from the father, who was not available for testing. Clearly the mother in this family had a different HHV-6B strain with a distinct DRR-pvT1 repeat pattern (Figure 7).

**Figure 7.**
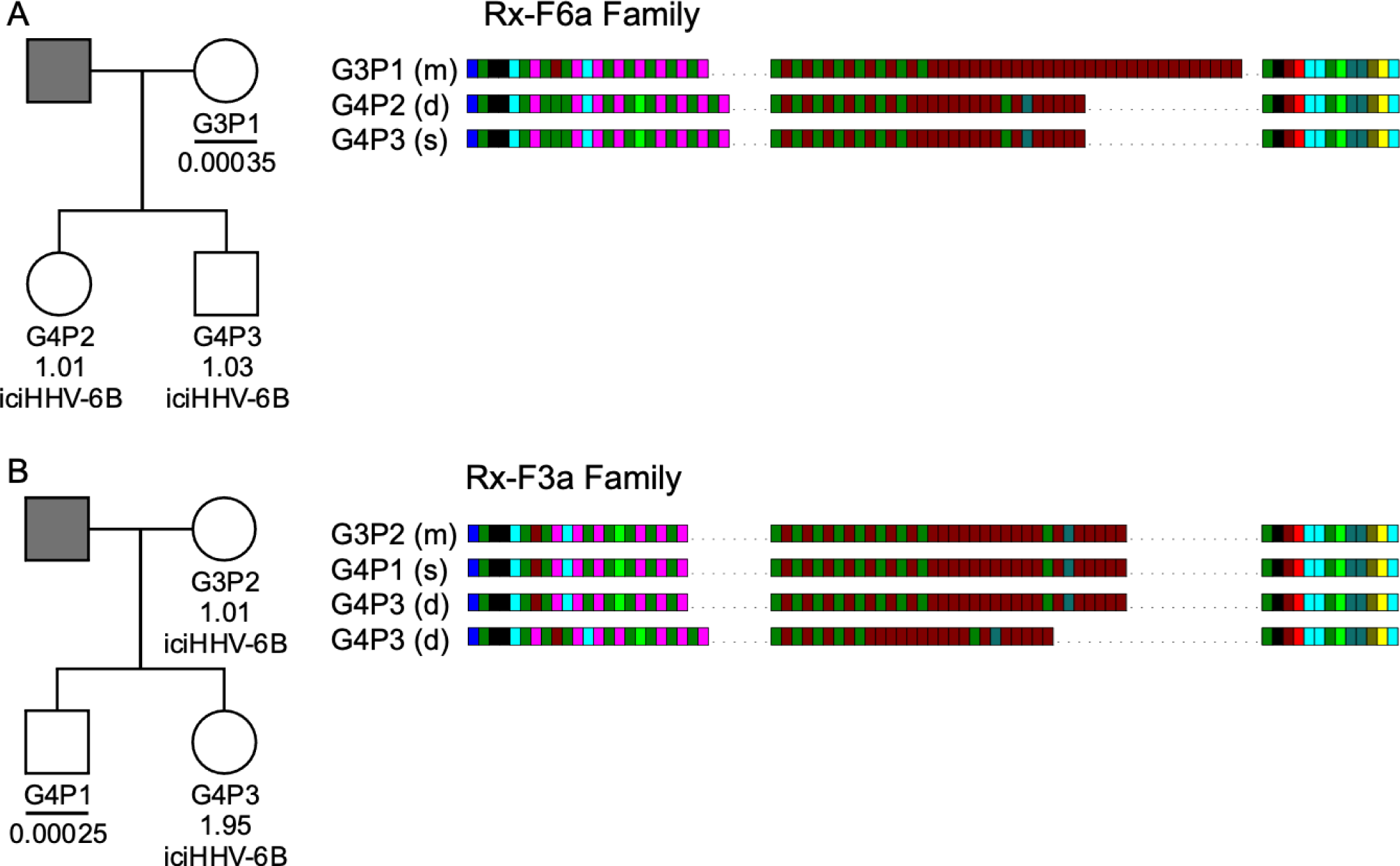
HHV-6B copy number and DRR-pvT1-repeat patterns can be used to identify potential iciHHV-6B reactivation and transmission within families. (A) In family Rx-F6a both children are iciHHV-6B carriers with approximately one copy per cell. They share the same DRR-pvT1 repeat map, presumably inherited from the father (grey filled square, not available for testing). The mother has a low level of acqHHV-6B in her saliva (0.00035 copies per cell) and the DRR-pvT1 repeat map is different from her children. (B) Evidence of iciHHV-6B reactivation in family Rx-F3a and transmission to non-iciHHV-6B son. Left shows the Rx-F3a family tree with HHV-6 copy number per cell in saliva DNA and iciHHV-6B carrier status. Father (grey filled square) was not available for testing. Right, the DRR-pvT1 repeat maps from family members. Daughter (G4P3) has two copies of iciHHV-6B, one copy shares the same DRR-pvT1 as seen in her mother (G3P2) and a second copy has a different DRR- pvT1 repeat pattern assumed to have been inherited from her father. The son (G4P1) has a very low level of HHV-6B in his saliva with the same pvT1 repeat pattern as the maternal iciHHV-6B genome. The DRR-pvT1 repeat patterns are also labeled with m (present in mother); d (present in daughter) and s (present in son).

Quantification of HHV-6B in saliva samples from another family (Rx-F3a) showed that the mother (G3P2) was an iciHHV-6B carrier and that her daughter (G4P3) had approximately 2 copies of iciHHV-6B per cell. Identical DRR-pvT1 repeat patterns between the mother and daughter proved maternal inheritance of one iciHHV-6B copy. A second, distinct DRR-pvT1 repeat pattern in the daughter was assumed to represent iciHHV-6B inherited from the father, who was not available for testing. The son (G4P1) had 0.00025 copies of HHV-6B per cell in saliva DNA, consistent with the level expected from acqHHV-6B infection. Importantly, the DRR-pvT1 repeat pattern in the son (G4P1) was identical to that in the mother and one copy of iciHHV-6B in his sister. This repeat pattern was not seen outside this family among 102 DRR-pvT1 repeat patterns (Figure 1 – figure supplement 1). These observations strongly suggest that iciHHV-6B genome excision occurred in the mother and that the reactivated HHV-6B was then transmitted to the non-iciHHV-6B son, who retained residual viral sequences in his saliva.

## Discussion

The discovery of germline transmission of chromosomally-integrated HHV-6A/6B genomes has raised questions about the possible relationship between iciHHV-6A/6B carrier status and lifelong disease risk (Gravel *et al*., 2015) (Gaccioli *et al*., 2020; Hill *et al*., 2017). Events that could have deleterious consequences include full viral reactivation (Endo *et al*., 2014; Gravel *et al*., 2013; Hall *et al*., 2010), intermittent expression of iciHHV-6A/6B genes that may elicit intermittent immune and inflammatory responses over a lifetime (Peddu *et al*., 2019), and an impact on telomere stability and function (Huang *et al*., 2014; Prusty *et al*., 2013; Wood & Royle, 2017; Zhang *et al*., 2017).

Progress towards understanding the potential impact of iciHHV-6A/6B carrier status has been advanced by the recent increase in the number of HHV-6A/6B genome sequences available (Aswad *et al*., 2021; Greninger *et al*., 2018; Zhang *et al*., 2017). This has shown that most iciHHV- 6A/6B genomes in populations are derived from a small number of ancient telomere-integration events. Here we have expanded the number of sequenced iciHHV-6A/6B genomes from individuals in Europe and North America and developed a web-based HHV-6 Explorer that can be used to interrogate diversity and functionality between inherited and acquired HHV-6A or HHV-6B genomes. The HHV-6 Explorer includes the sequence of 84 complete or nearly complete iciHHV-6A/6B genomes representative of the various known clades in the phylogenetic network and genome sequences from 28 HHV-6A/6B viruses circulating in populations. Using the HHV-6 Explorer we displayed two mutations that introduced stop codons in U79 in three of 14 iciHHV-6A samples in the 17p clade and in U14 in one of three iciHHV-6B samples in the 17p (minor) clade (Figure 2 – figure supplement 1) so demonstrating that each mutation arose after integration. The presence of an inactivating iciHHV-6A/6B mutation in some but not all descendants of a particular ancestral integration event adds a further complication to understanding the potential lifetime risk for individual iciHHV-6A/6B carriers.

The relationship between iciHHV-6A/6B carrier status and potential associated disease risk may also be influenced by interactions between the integrated viral genome and genes or chromatin on the chromosome carrying the viral genome (Gravel *et al*., 2015; Wood & Royle, 2017). It is therefore important to have a rapid method to distinguish between viral strains and integration sites, which is offered by the hypervariable pvT1 region in HHV-6B. Length variation across pvT1 is mostly attributable to the diverse interspersion of (CTAGGG) and canonical (TTAGGG) telomere repeat sequences in the middle of pvT1 amplicons. Notably, short blocks of (CTAGGG)n repeats are highly unstable in human telomeres in somatic and germline cells, probably due to replication errors and *in vitro* studies showed that (CTAGGG)n repeats bind efficiently to POT1, a component of Shelterin (Barrientos *et al*, 2008; Baumann & Cech, 2001; de Lange, 2018; Lim *et al*, 2009; Loayza *et al*, 2004; Mendez-Bermudez *et al*, 2009). In many cases analysis of the DRR-pvT1 repeat pattern alone can be used as a rapid and inexpensive way to predict the phylogenetic clade and integration site of an iciHHV-6B genome.

DRR-pvT1 analysis is also an excellent tool for tracking acqHHV-6B transmission in families as shown by the evidence that community-acquired HHV-6 transmission usually occurs between family members, with maternal transmission most common. We showed that DRR-pvT1 repeat patterns can be used effectively to discriminate between HHV-6B strains circulating in communities (61/63 different pvT1 sequences in saliva from healthy non-iciHHV-6B donors in the UK). In the future, pvT1 analysis could be used to trace patterns of transmission more generally. This may be particularly important in the setting of organ and tissue transplants (Hill, 2019). For example, reactivation of iciHHV-6B from a donor tissue could be monitored and differentiated from HHV-6B acquired by the recipient in early childhood or to identify cases of multiple infections by different strains of HHV-6B.

The iciHHV-6B and free HHV-6B viruses in communities must have evolutionary histories that are interlaced but these are difficult to disentangle as there is little understanding of HHV-6 telomeric integration and iciHHV-6B excision and reactivation. The current picture indicates a modest number of ancient iciHHV-6B clades (in Europe and North America at least) some of which are accumulating mutations that would prevent full reactivation (discussed above). There is also at least one example of an iciHHV-6B genome with high sequence homology to a group of genome sequences from acqHHV-6B (Aswad *et al*., 2021; Greninger *et al*., 2018), which complicates interpretation of HHV-6B phylogenetic relationships (Forni *et al*, 2020). Indirectly this suggests that some modern circulating strains retain the capacity for germline integration into telomeres and to be passed on or that iciHHV-6B can reactive and be transmitted, re-entering the reservoir of circulating strains. We demonstrated that some copies of HHV-6B in saliva acquire a telomere, indicative of integration in somatic cells *in vivo* and as previously detected *in vitro* (Arbuckle *et al*., 2010). The viral load in the saliva samples was low, as seen in some other studies (Leibovitch *et al*, 2014; Leibovitch *et al*, 2019; Turriziani *et al*, 2014), and the number of HHV-6B genomes with a telomere was small (Figure 4, Supplementary Table 3), which is consistent with the majority of HHV- 6B genomes present as viral particles in saliva (Jarrett *et al*, 1990). The approach we used to detect telomeric integration in saliva could, in principle, be used to detect HHV-6B integration events in sperm from healthy non-iciHHV-6B men (Neofytou *et al*, 2009) (Godet *et al*, 2015; Kaspersen *et al*, 2012). This would address the outstanding question of whether current HHV-6B strains circulating in communities can integrate in the germline.

Previously we have proposed that iciHHV6A/6B genomes can be excised from telomeres in a one or two-step process as by-products of t-loop processing (Huang *et al*., 2014; Wood & Royle, 2017). In this study we measured the frequency of iciHHV-6B excision events, by detection of novel telomeres at DRL-T2 and for the first time at DRR-T1, in unrelated iciHHV-6B carriers and in different tissues and cell types. The frequencies of iciHHV-6B truncations at DRL-T2 (range, 0.014 - 0.0031 per cell) and DRR-T1 (range, 0.0002-0.0029 per cell) are not directly comparable because they were ascertained by different approaches but both are surprisingly high *in vivo* (Figure 5, Figure 6). This demonstrates that tissues in iciHHV-6B carriers must be mosaic for different compositions of the iciHHV-6B genome. The telomeres formed at DRL-T2 were expected to be particularly short, limited to the length of the (TTAGGG) repeat array at T2. However, a cell with a small number of very short telomeres will elicit a telomere-mediated DNA damage response and entry into cellular senescence (Cesare *et al*, 2013; d’Adda di Fagagna *et al*, 2003; de Lange, 2018; Takai *et al*, 2003) so preventing proliferation of a cell with a potentially unstable residual iciHHV-6B genome. It is therefore remarkable that a high percentage of newly formed telomeres at DRL-T2 were lengthened. The frequency of DRL-T2 telomere lengthening was highest in circulating white blood cells and in two pluripotent cell lines (Figure 5) consistent with lengthening by telomerase. From these observations and the increase in viral genome excision events in TSA treated cells, we also propose that iciHHV- 6A/6B genomes are more likely to be excised and reactivated in undifferentiated or pluripotent cells. By extrapolation, the potential for iciHHV-6A/6B genome excision, viral gene expression or full reactivation has the potential to be deleterious during early development (Miura *et al*., 2020).

Finally, we present two lines of evidence that iciHHV-6B genome can be excised and re-enter the reservoir of circulating HHV-6B strains. First, we identified individuals with full-length iciHHV-6B or DR-only genomes that fall into the same clade in the phylogenetic tree (Figure 3). These integrated full-length iciHHV-6B or DR-only genomes share similar pvT1 repeat patterns and subterminal-iciHHV-6B junction sequences. This demonstrates that the different iciHHV-6B structures are carried in the same telomere-allele at the particular chromosome end and it proves that these DR-only carriers have arisen by germline excision of U-DR in an ancestor. The fact that this has occurred independently at least twice suggests this is a relatively common event, and this is supported by our evidence that new telomere formation at DRR-T1 occurs at a measurable frequency in sperm DNA. Second, we show an example of probable reactivation of iciHHV-6B in a mother with transmission to her son, who carries a very low load of acqHHV-6B with the same distinctive DRR-pvT1 repeat pattern in his saliva. This observation warrants further research to determine how frequently HHV-6B is transmitted from iciHHV-6B parent to their non-iciHHV-6B children but it requires the use of a hypervariable marker, such as pvT1, that has the power to discriminate between HHV-6B strains.

## Materials and Methods

### Saliva collection and DNA extraction

The study was conducted in accordance with the Declaration of Helsinki and with the approval of the University of Leicester’s Research Ethics Committee (refs: 10553-njr-genetics; njr-61d3). Saliva samples were donated by individuals and members of families (all 18 years or older) with informed consent and were given anonymous identifiers at the point of collection (Garrido-Navas *et al*, 2020). Approximately 1.5 ml of saliva was collected in a OraGEN saliva collection tube (Genotek, Ottawa, Canada), and DNA was extracted from 500 µl following the manufacturer’s instructions.

### Identification of DNA samples and cell lines with iciHHV-6A/6B or acqHHV-6B

The full list of iciHHV-6A/6B-positive DNA samples and cell lines used in this study is given in Supplementary Table 1 with available information on donor ethnicity and iciHHV-6A/6B chromosomal location. To interrogate any relationship between iciHHV-6A/6B carrier status and heart failure, the BIOlogy Study to TAilored Treatment in Chronic Heart Failure (BIOSTAT-CHF) cohort (Voors *et al*, 2016) was screened. The BIOSTAT-CHF study complied with the Declaration of Helsinki and was approved by the relevant ethics committee in each centre, while all participants gave their written, informed consent to participate. The 2470 blood DNAs from Europeans in the BIOSTAT-CHF cohort was screened using PCR assays to detect various regions of the viral genome (U11, DR3 and DR5 in HHV-6A, and DR7 in HHV-6B) (Huang *et al*., 2014; Zhang *et al*., 2017). This identified 19 iciHHV-6A/6B positive samples (nine iciHHV-6A, 0.36%, and ten iciHHV-6B, 0.40%, one of which was iciHHV-6B DR-only). Four iciHHV-6B carriers were identified among the saliva donors: two with a single copy of iciHHV-6B, one with two copies of iciHHV-6B, and one with iciHHV-6B DR-only. The saliva samples were also screened for acqHHV-6B using ddPCR, described below. Additionally, two kidney DNA samples (part of the TRANScriptome of renaL humAn TissuE (TRANSLATE) study (Marques *et al*, 2015; Rowland *et al*, 2019)) were positive for DR3 by PCR assay and were confirmed as acqHHV-6B by ddPCR.

### HHV-6B quantification by ddPCR

HHV-6B copy number was estimated by ddPCR as describe previously (Bell *et al*, 2014). Hydrolysis probe ddPCRs (20 μl volumes) consisted of 1 x ddPCR Supermix for probes without dUTP (Bio-Rad Laboratories), virus-specific primers (HHV-6B POL F and HHV-6B POL R) and FAM-labelled HHV-6B POL (U38) probe (Eurogentec; Supplementary Table 6) at 300 nM and 200 nM respectively, 1 x RPP30 primer/HEX labelled probe mix (Bio-Rad Laboratories), 1μl XhoI digestion mix consisting of 5U XhoI restriction enzyme and 1 x NEB buffer 2.1 (New England Biolabs), and 10 ng genomic DNA for iciHHV-6B samples or 200 ng DNA for non-iciHHV-6B samples. Assays to quantify acqHHV-6B were carried out in triplicate. DNA from the HT1080 osteosarcoma-derived, established cell line was used as a negative control to detect non-specific amplification or contamination, and water was used as a no-template control. As very low viral copy number were expected, the ddPCR reactions were set up in a PCR-clean room where no HHV-6 PCR had previously been conducted. Droplets were generated using QX200 Droplet Generator (Bio-Rad Laboratories) with 70 μl Droplet Generation Oil for Probes or 70 μl Droplet Generation Oil for Evegreen, as appropriate (Bio-Rad Laboratories). Thermocycling was carried out according to the manufacturer’s instructions on a Veriti (Bio-Rad Laboratories). QX200 Droplet Reader and QuantaSoft analysis software (Bio-Rad Laboratories) were used to count droplets and to calculate the estimated copy number. The number of DR copies per cell in iciHHV-6B DNA was determine by ddPCR with primers DR6B-F and DR6B-R and DR6B FAM-labelled hydrolysis probe (Eurogentec, Liège, Belgium) together with the HEX-labelled RPP30 reference probe (Bell *et al*., 2014).

Absolute quantification of DR and PAC1 was determined by ddPCR using DNA intercalating dye, EvaGreen. EvaGreen ddPCRs (20 μl volumes) consisted of 1 x ddPCR EvaGreen Supermix (Bio-Rad Laboratories), virus-specific primers for DR (DR6B-F and DR6B-R) or PAC1 (PAC1F and PAC1R-33) at 300 nM, 1μl XhoI digestion mix consisting of 5U XhoI restriction enzyme and 1 x NEB buffer 2.1 (New England Biolabs, Ipswich, Massachusetts), and 10 ng genomic DNA for iciHHV-6B samples or 200 ng DNA for non-iciHHV-6B samples.

### Cell culture and TSA treatment

LCLs with iciHHV-6B were cultured in RPMI 1640 medium as described previously (Huang *et al*., 2014). The CRL-1730 adherent human vascular endothelial cell line derived from umbilical cord vein (ATCC, (Shioda *et al*., 2018)) was cultured in F-12K medium (Gibco/ThermoFisher Scientific) supplemented with 10% heat inactivated foetal bovine serum (Gibco/ThermoFisher Scientific), 0.1 mg/ml heparin (Sigma-Aldrich, Darmstadt, Germany) and 1% endothelial growth supplement (BD Biosciences). The adherent mesenchymal stem cell line derived by explant culture from umblical cord, d37 (gifted by Dr Sukhvir Rai, University of Leicester), was cultured in human mesenchymal stem cell growth media (MSCBM hMSC Basal Medium, Lonza, Basel, Switzerland) supplemented with mesenchymal stem cell growth supplements (MSCGM hMSC SingleQuot Kit, Lonza, Basel, Switzerland).

For TSA treatment, 2 x 10^6^ cells from the 4B-11p15.5 iciHHV-6B LCL were suspended in RPMI 1640 medium supplemented with 0, 100 or 200 ng/ml Trichostatin A (Sigma-Aldrich, Darmstadt, Germany) in three biological replicates. After 124 h (5 days), cells were pelleted at 1200 rpm for 8 min, washed twice with phosphate-buffered saline and snap-frozen on dry ice. DNA was extracted using phenol-chloroform and precipitated using ethanol. The DNA pellet was washed with 80% ethanol, briefly air-dried and dissolved in nuclease-free water. DRL-T2 STELA (see below) was conducted on extracted DNA using primer UDL6R to detect telomeres at DRL T2, as described previously (Huang *et al*.). STELA was carried out two times on DNA from each treatment from three biological replicates.

### Whole iciHHV-6A/6B genome sequencing

Ten iciHHV-6A and five iciHHV-6B genomes were sequenced and annotated as described previously (Huang *et al*., 2014; Zhang *et al*., 2017). Pooled, overlapping PCR amplicons (100 ng) were sheared using a Covaris S220 sonicator to an approximate size of 450 bp. Sequencing libraries were prepared using the LTP library preparation kit for Illumina platforms (Kapa Biosystems, Germany). The libraries were then processed for PCR (7 cycles) using the LTP library preparation kit and employing NEBNext multiplex oligos for Illumina index primer pairs set 1 (New England Biolabs, Ipswich, Massachusetts). Sequencing was performed using a NextSeq 500/550 mid-output v2.5 300 cycle cartridge (Illumina, San Diego, California) to produce 6-9 million paired-end 150 base reads per sample. The annotated sequences were deposited in NCBI GenBank under accession numbers: MW049313-MW049327.

### Genome sequence analysis

94 complete or near-complete HHV-6A/6B sequences were downloaded from NCBI GenBank (Supplementary Table 1) and, together with the 15 newly sequenced genomes, were aligned using MAFFT (v7.407, (Katoh & Standley, 2013)) with default parameters. The alignment was trimmed using trimAI (v1.4.1, (Capella-Gutierrez *et al*, 2009)) to remove gaps. Phylogenetic networks were inferred using FastME (v2.1.6.1), a distance-based algorithm, with bootstrap values calculated from 100 replicates. Phylogenetic networks were plotted using Interactive Tree of Life browser application (https://itol.embl.de, v6.1) (Letunic & Bork, 2016). The time to the most recent common ancestor was estimated using Network 10.0 (Fluxus-engineering, Colchester, United Kingdom) using rho values calculated by Network (Forster *et al*, 1996) and an assumed mutation rate of 0.5 x 10^-9^ substitutions per base per year (Scally & Durbin, 2012; Zhang *et al*., 2017).

### Development of the HHV-6 explorer

The HHV-6 Counter takes a fasta multiple alignment and compares each sequence to a selected reference HHV-6 strain to generate counts of genetic variation (substitutions, insertions, deletions) from overlapping or non-overlapping windows across the HHV-6 genome. If a corresponding genbank file for the reference is present, HHV-6 Counter will also provide windowed counts for coding sequence and amino acid variation across the genome and for each gene. This includes missense/nonsense changes, in-frame insertions/deletions, nonsense insertions, potential splice site changes, loss of start or stop and tracking of frameshift changes. Windows which contain non-standard base characters or have sequence gaps due to method failures are flagged but their counts are still retained in the results. HHV-6 Counter exports the count windows both as a series of Excel files and as a python panda pickle file for use in the HHV-6 Explorer.

The HHV-6 Explorer is based on plotly Dash (https://dash.plotly.com/) and, using the output from HHV-6 Counter, allows the graphical display of variation counts for different strains across the HHV-6 genomes and/or on a per gene basis compared against a selected reference HHV-6 strain. It also displays a multiple alignment for the selected gene. A pre-populated version of HHV-6 Explorer containing the data from this manuscript can be found at https://www.hhv6explorer.org/. The source code for HHV-6 Explorer and HHV-6 Counter is available from https://github.com/colinveal/HHV6-Explorer.

### PCR primers and other oligonucleotides

The primer sequences used in this study and the Taqman hydrolysis probes are listed in Supplementary Table 6.

### Amplification and analysis of subtelomere-iciHHV-6B junctions

Subtelomere-iciHHV-6B junctions were amplified by PCR (33 cycles) using a trial an error approach with primer DR8F(A/B) and various primers known to anneal to chromosomal subtelomere regions (Zhang *et al*., 2017). Amplicons were purified using a Zymoclean gel DNA recovery kit (Zymo Research, Irvine, California) and sequenced using primer DR8FT2 or the appropriate subtelomere primer (Supplementary Table 6). The number and pattern of telomere repeats (TTAGGG) and degenerate telomere-like repeats across the subtelomere-iciHHV-6B junction amplicons were identified and colour-coded manually to generate repeat patterns.

### Amplification and analysis of HHV-6B DRR-pvT1 and DRL-pvT1

DRR- pvT1 was amplified by PCR using primers DR1R and U100Fw2 in the first round (94°C, 1.5 min; 25 cycles 94°C 15 s, 62°C 30 s, 68°C 10 min; 68°C 2 min) and primers DR421R and TJ1F in the second round (94°C, 1.5 min; 25 cycles 94°C 15 s, 64°C 30 s, 68°C 1.5 min; 68°C 2 min). DRL-pvT1 was amplified by STELA (see below) followed by a secondary PCR using primers DR421R and TJ1F. The short pvT1 amplicons were size-separated by electrophoresis in a 3% NuSieve (Lonza, Basel, Switzerland) agarose gel, extracted and Sanger sequenced using primer TJ1F. The number and pattern of (TTAGGG) and degenerate telomere-like repeats were identified and colour-coded manually to generate repeat maps.

### STELA and detection of newly formed telomeres at DRR-T1 and DRL-T2

The telomere at the end of the iciHHV-6B genome was amplified by STELA using the DR1R primer and Telorette2/Teltail essentially as described previously (Huang *et al*., 2014) (Jeyapalan *et al*, 2008). DNA was diluted to 250-1000 pg/µl for cell line DNA, 500 pg/µl for saliva DNA, 600 pg/µl for blood DNA and 1000 pg/µl for sperm DNA. The primer concentrations in each 10 µl STELA reaction were 0.3 µM DR1R, 0.225 µM Telorette2 and 0.05 µM Teltail. *Taq* polymerase (Kapa Biosystems, Germany) was used at 0.04 U/µl and *Pwo* (Genaxxon Bioscience, Ulm, Germany) at 0.025 U/µl. The STELA PCRs were cycled 25 times.

To detect telomeres at DRR-T1, STELA was conducted as above using primers DR1R and Telorette2/Teltail on iciHHV-6B DNA samples that showed a length difference between DRL-pvT1 and DRR-pvT1. Next the STELA reaction product (1µl) was diluted 1:10 in water and used as input for PCR of pvT1 using primers DR421R and TJ1F (25 cycles). The amplicons were size-separated by agarose gel electrophoresis to distinguish DRR-T1-associated telomeres from the majority of DRL-T1- associated telomeres. The remainder of the undiluted STELA product was size-separated by agarose gel electrophoresis and amplified telomeres detected by Southern blot hybridisation to a radiolabelled (TTAGGG)n probe.

To detect telomeres at DRL-T2, primer UDL6R was used in STELA reactions instead of primer DR1R with 250-1000 pg genomic DNA per reaction and cycled 26 times. Amplicons hybridising to the radiolabelled (TTAGGG)n probe that migrated at less than 900 bp were counted as unlengthened truncations, those between 900 bp and 8.6 kb were counted as lengthened truncations, and those larger than 8.6 kb were not counted as truncations. The number of truncations per cell was estimated by converting the amount of input DNA to cell equivalents on the assumption that a cell contains 6.6 pg DNA (or 3.3 pg for a haploid sperm cell) and dividing the number of truncations (lengthened and unlengthened) by the number of cells screened. To sequence DRL-T2-associated telomeres, STELA products were extracted from agarose gel slices using repeated freeze-thawing and reamplified using primers DR8RT2 and Telorette2. The recovered amplicons were Sanger sequenced.

### Detection of integrated HHV-6B in individuals with acqHHV-6B

STELA was carried out on genomic DNA from individuals with a known copy number of acqHHV-6B, measured by ddPCR. As the copy number of acqHHV-6B was low and telomere integration events expected to be lower, various precautions were introduced for these experiments. STELA reactions were set up in a room previously unused for any STELA or HHV-6 experiments. In addition to avoid contamination with previously generated iciHHV-6A/6B STELA products, STELA primers with new barcodes (Telorette2BC28/Teltail2BC38) and a new a flanking primer (DR2RSTELA) were used. The DR2RSTELA primer anneals upstream of the DR1R primer used for other iciHHV-6A/6B STELA reactions. STELA reactions were set up as explained above, but with 2-5 ng genomic DNA per reaction and cycles 25 times. Next 1 µl of STELA reaction product was diluted 1:10 in water and used as input for amplification of DRL-pvT1, using primers DR421R and TJ1F (25 cycles). The amplicons were sequenced using TJ1F to confirm that the DRL-pvT1 sequence in the amplified telomere was the same HHV-6B strain as present in that saliva sample. The remainder of the undiluted STELA product was size-separated by agarose gel electrophoresis and detected by Southern blot hybridisation to a radiolabelled (TTAGGG)n probe. As a control for non-specific amplification from another telomere or elsewhere in the human genome, 900 STELA reactions were conducted each with 5 ng genomic DNA from a HHV-6A/6B free cell line, HT1080 (4500 ng total). No STELA amplicons were generated from the HT1080 control reactions.

### DR circles

Double restriction digests were carried out at 37C for 1 hour using 10 U of each enzyme (XbaI and ScaI-HF, or PstI and SacI) and 1x NEB CutSmart Buffer (New England Biolabs, Ipswich, Massachusetts) in 500 uL reactions containing 10 μg 4B-11p15.5 or 1B-HHV-6B DNA. Enzymes were heat inactivated at 80C for 20 minutes. Control DNA was treated without any restriction enzymes. Diluted DNA was amplified using PCR with primers DR8F(A/B) and DR3R with a 10 minute extension time (26 cycles). PCR products were size-separated on a 0.8% agarose gel with electrophoresis and amplicons were detected by Southern blot hybridisation to a radiolabelled telomere (TTAGGG)n probe.

### Statistical analysis

Data are expressed as means ± standard error of the mean (SEM). Mann-Whitney test was used to compare unpaired groups of ranked data obtained from assays conducted on DNA derived from the same biological sample type (Figure 4C, 4F). The Wilcoxon test was used to compare paired groups of data obtained from treated and untreated samples. Statistical analyses were performed using Prism software (Version 9.0, Graphpad Software). Outliers were not removed from data sets. Samples sizes were based on availability of suitable biological materials.

## Acknowledgements

We thank Dr Sukhvir Rai (University of Leicester) for the gift of the d37 mesenchymal stem cell line; Dr Yan Huang, Dr Enjie Zhang, Dilan Patel and Ryan Mate for preliminary work that contributed the initiation of this study; and Drs Chiara Batini, Celia May and Jon Wetton for their advice on phylogenetics and hypervariable markers. We also thank Matthew Denniff and Charlotte Hogg for help with sample acquisition and TRAP assays respectively. We also acknowledge the contribution of members of the BIOSTAT-CHF consortium.

## Figure Supplements and Supplementary Tables

**Figure 1 – figure supplement 1.**
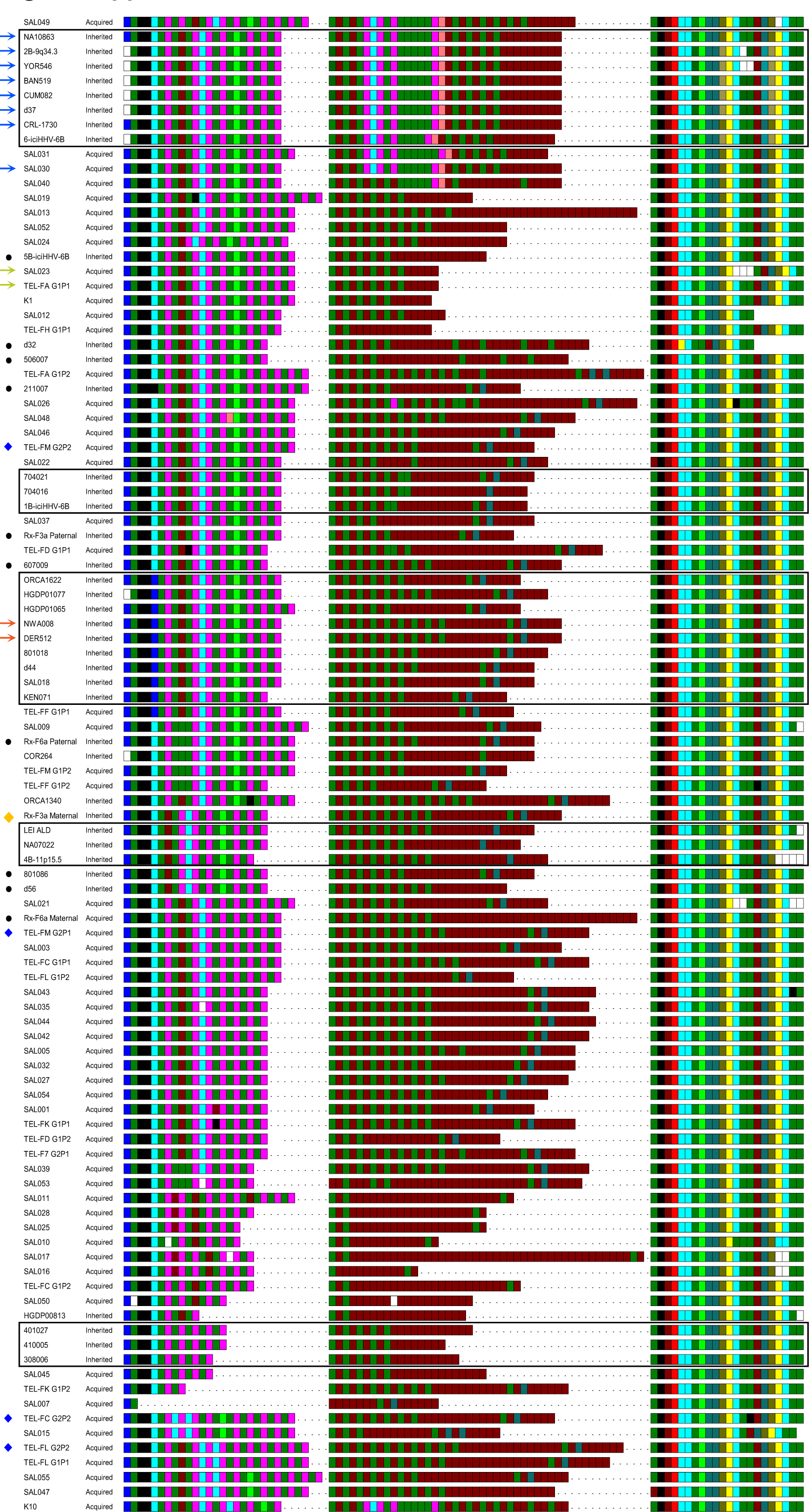
Complete set of pvT1 interspersion maps from DR_R_ in iciHHV-6B (inherited) and acquired HHV-6B (non-inherited) strains. The 102 DR_R_-pvT1 repeat patterns shown demonstrate the diversity among iciHHV-6B (inherited) and HHV-6B (acquired) genomes. Ninety of the repeat patterns were unique. Degenerate, telomere-like repeats present in the HHV-6B pvT1 region are colour-coded: Dark green, TTAGGG; brown, CTAGGG; cyan, TTAGTG; yellow, TTACTG; dark yellow, ATAGAC; teal, CTAAGG; pink, CTATGG; lime green, TTATGG; blue, GTAGTG; peach, TTAGAG; red, GTCTGG. Black squares represent other, less common degenerate repeats and white squares show where the sequence could not be determined accurately. Dashes between repeats were added to maximize alignment between samples allowing comparison between the left, middle (highly variable) and right (highly conserved) sections of DR_R_-pvT1. Repeat maps were grouped primarily based on the left section, then on less common features within the middle or right region, and finally by the length of the middle region. Ninety eight of the 102 DR_R_-pvT1 repeat patterns are from unrelated individuals and four are from children who have an acqHHV- 6B that has a different DR_R_-pvT1 repeat pattern from their parents (blue diamonds). Coloured arrows identify 12 repeats patterns found more than once in donors not known to be related. Among eight of the identical DR_R_-pvT1 repeat patterns (blue arrows), seven are in iciHHV-6B genomes predicted to be integrated in the 9q telomere and one in an acquired HHV-6B in SAL030. Two others, in NWA008 and DER512 from the UK, have a 17p iciHHV-6B integration and share identical DR_R_-pvT1 repeat patterns. In addition, the acquired HHV-6B strains in SAL023 and TEL-FA G1P1 share the same DR_R_-pvT1 repeat pattern. Black boxes surround repeat maps from iciHHV-6Bs predicted to have the same integration site, by whole viral genome sequence homology, FISH, or subtelomere-iciHHV-6B junction sequence. Notably within integration groups there are often shared characteristic features in the DR_R_-pvT1 repeat patterns. Black dots indicate iciHHV-6B samples where DR_R_-pvT1 could not be used to predict the integration site confidently. For example, the DR_R_-pvT1 in iciHHV-6B in d32 has a unique right section and is unlikely to share common ancestry with any of the other iciHHV-6B carriers analysed. Yellow diamond identifies the DR_R_-pvT1 repeat pattern in family Rx-F3a, transmitted from iciHHV-6B mother to non-carrier son. This pvT1 repeat map is different from the other 101 maps shown here.

**Figure 1 – figure supplement 2.**
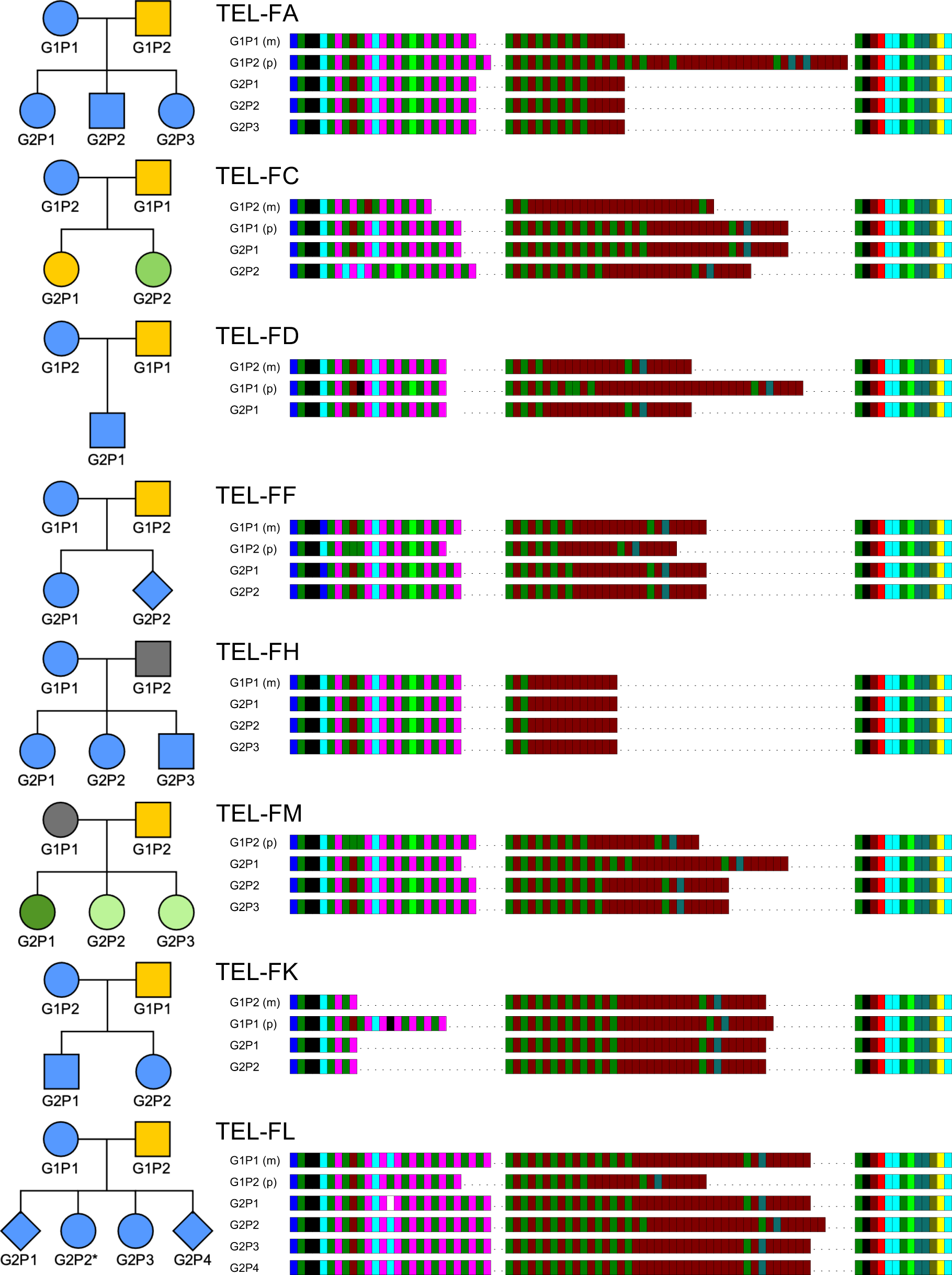
HHV-6B strain identification using DR_R_-pvT1 repeat patterns in eight families, suggests that transmission of acqHHV-6B is predominantly from parents. The key to the repeat pattern is the same as in Figure 1. Pedigree symbols shaded blue indicate the child(ren) had the same acqHHV-6B DR_R_-pvT1 repeat pattern as their mother and yellow indicates it was the same as their father. The pedigree symbols shaded grey identify individuals from whom the pvT1 region could not be amplified. The green symbols identify children that do not share a DR_R_-pvT1 map with either parent and the dark green symbol for TEL-FM G2P1 indicates that this individual had a DR_R_- pvT1 that was different to their siblings and parents. TEL-FL G2P2 (asterisk) has a DR_R_-pvT1 repeat pattern that differs from that in their mother by a gain of two repeats (CTAGGG-TTAGGG) in the middle section.

**Figure 2 – figure supplement 1.**
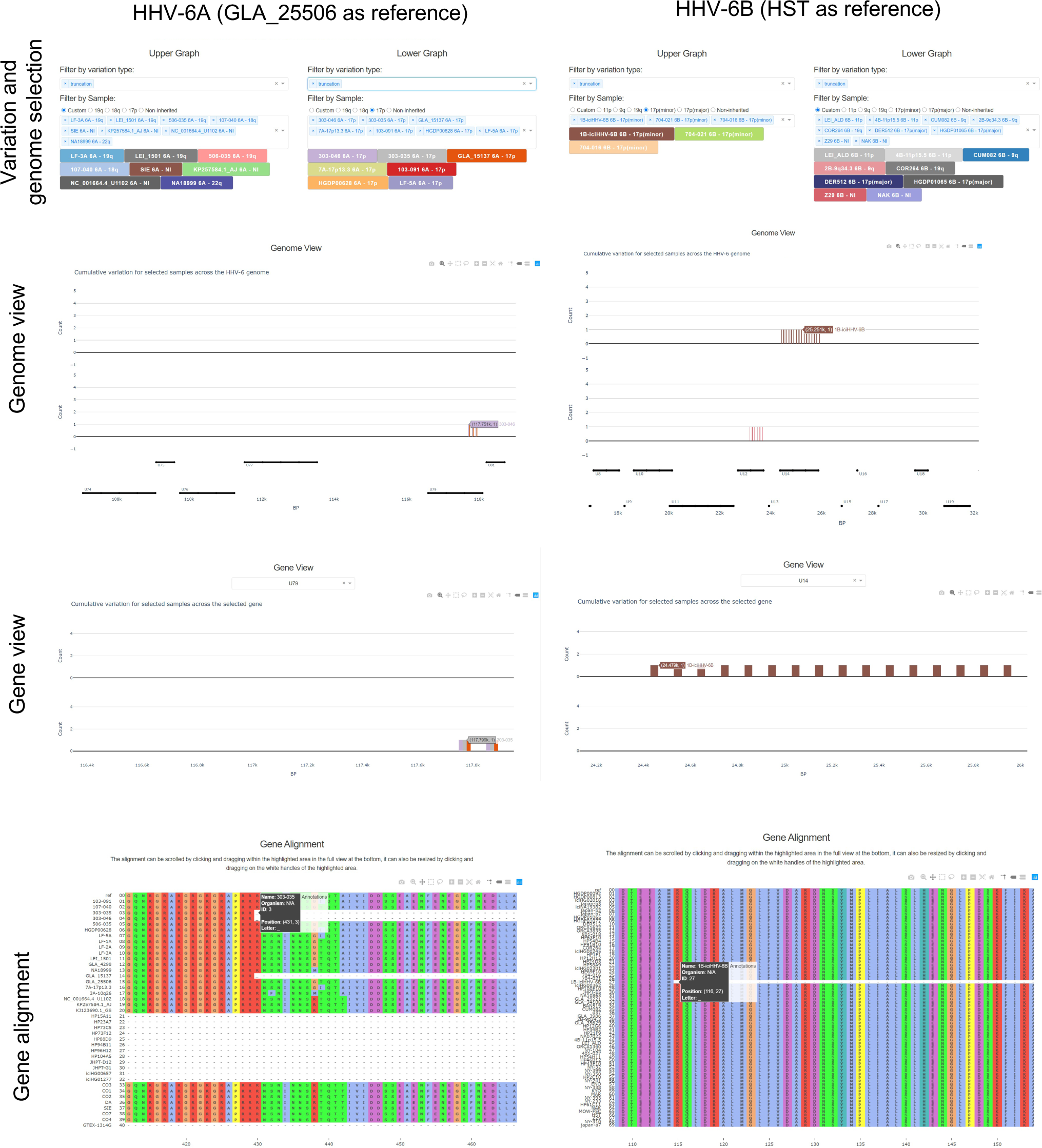
Examples of views taken from the HHV-6 Explorer. The HHV-6 explorer allows different types of variation across iciHHV-6A/6B and acquired HHV-6A/6B genomes from different individuals to be compared against a reference genome. The user can select the size of overlapping or non-overlapping sliding windows in an upper and lower graph to allow easier comparison when a larger number of genomes are selected from the drop-down menus. After selecting the type(s) of variation and genomes to be explored, a lower and upper Genome View graph will be generated based on the user-selected sliding window size. The Genome View shows the specified type(s) of variation across the whole genome as a count of mutations per window size (100 bp, non-overlapping in the examples shown here). Cumulative mutations per window size are shown if multiple types of variation are selected. Each genome is assigned a unique colour and where part of sequence is missing, a value of negative one will be assigned for that particular window and the bar on the graph will be red. In the examples shown, iciHHV-6A and acquired HHV-6A genomes have been compared to GLA_25506 (19q iciHHV-6A) as the reference, and iciHHV-6B and acquired HHV-6B genomes have been compared to HST (acquired HHV-6B). In both examples, truncations (i.e. a nonsense mutation encoding a premature stop codon) are displayed. For HHV-6A, seven 17p iciHHV-6A genomes have been selected to be displayed on the lower graph and a variety of acquired HHV-6A and 18q, 19q and 22q iciHHV-6A genomes have been selected to be displayed on the upper graph. The Genome View is zoomed in to show a region displaying truncations in U79 in three 17p iciHHV-6A genomes (303-046, 303-035 and GLA_15137). In the HHV-6 Explorer the next graph generated is the Gene View where a single gene of interest can be selected to view variation at the amino acid level, U79 in this case. The final plot on the HHV-6 Explorer is the Gene Alignment, based on an alignment of all iciHHV-6A and acquired HHV-6A (or iciHHV-6B and acquired HHV-6B) used to generate the phylogenetic networks. In this region of U79, the sequence assemblies are missing data in 13 of the viral genomes (dashes). The exact amino acid position of the premature stop codon for 303-035 is highlighted, underscore at amino acid 431. All other iciHHV-6A/HHV-6A genomes (for which sequence is available) lack this premature stop codon, including other 17p iciHHV-6A genomes (e.g. 7A-17p13.3) indicating that this mutation arose after integration. In the right-hand column, members of the iciHHV-6B 17p (minor) integration group are displayed in the upper graphs and a variety of other iciHHV-6B integrations and acquired HHV- 6B genomes are displayed on the lower graphs. HST was used as the reference strain here. A premature stop codon is present at position 116 in U14 in 1B-iciHHV-6B but not in 704-021 or 704-016, which share the same common ancestor. Again, this indicates that the nonsense mutation arose after integration.

**Figure 5 – figure supplement 1.**
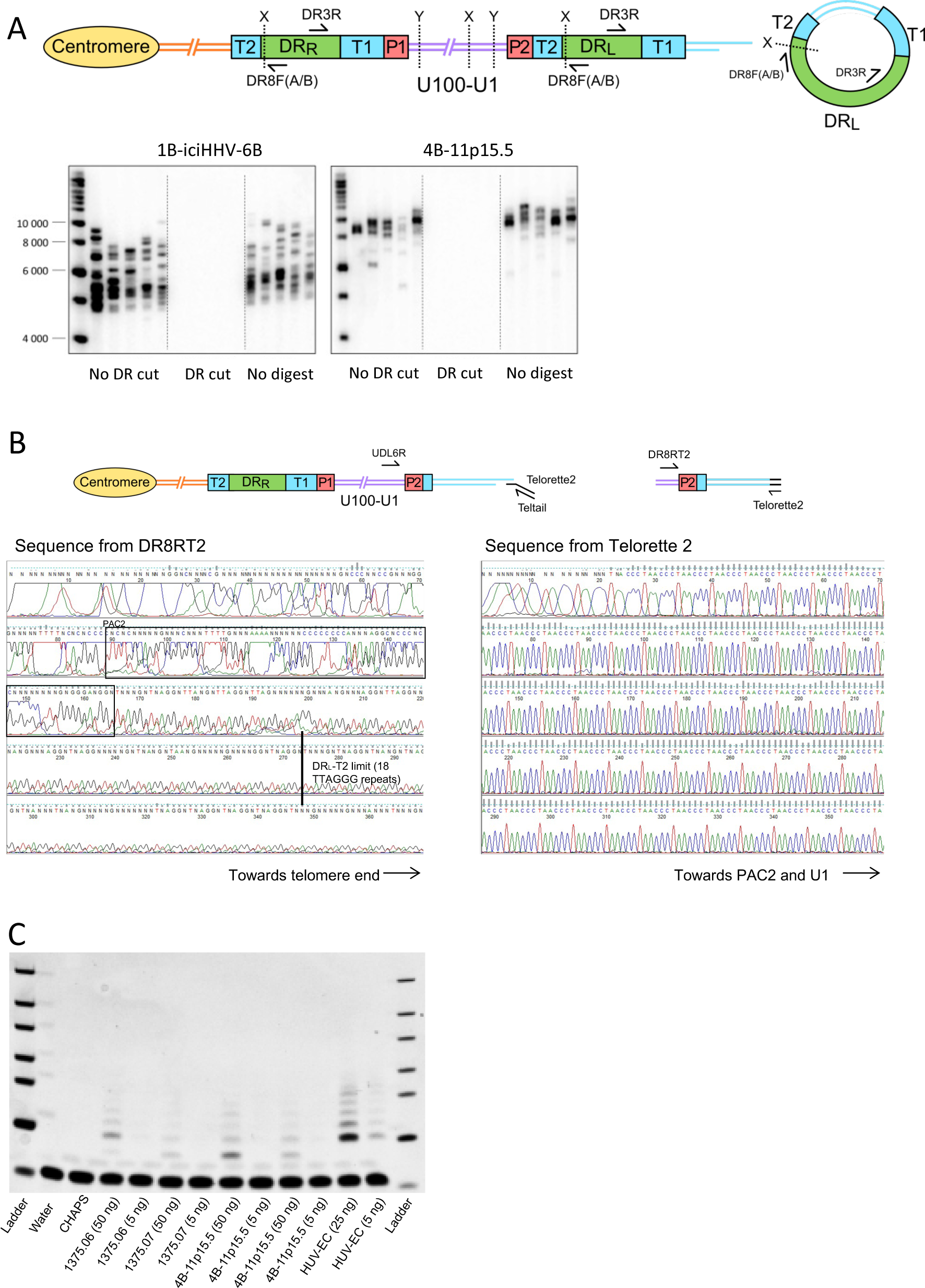
Detect of excised circular DR-only molecules, sequencing of a lengthened new telomere at DR_L_-T2 and detection of telomerase activity. (A) Schematic of DR-circle assay. DR-circles were amplified using primers (DR8F(A/B) and DR3R) that should not generate a PCR product from full-length iciHHV-6B but should amplify across telomere repeats in a DR-circle, the predicted reciprocal product of a t-loop mediated excision event at DR_L_-T2. Genomic DNA from two iciHHV-6B individuals (1B-iciHHV-6B and 4B-11p15.5) was digested with combinations of restriction enzymes that cut in the unique region and DR (X: SacI and PstI), only the unique region (Y: XbaI and ScaHF) or were not treated with restriction enzymes. Treated DNA was amplified using DR8F(A/B) and DR3R, and PCR products were size separated and detected by Southern blot hybridized to a radiolabelled (TTAGGG)_n_ telomere probe. Amplicons of variable length were detected reflecting different lengths of telomere repeat arrays expected in individual DR-circles. Restriction digestion with SacI/PstI, including between DR8F(A/B) and T2, prevents amplification from DR-circles. Importantly, the minimal size of amplicons detected was greater than the combined length of the flanking regions. Full length 4B-11p15.5 had longer average telomere length at DR_L_-T1 than 1B-iciHHV-6B, which is consistent with the longer products generated in the DR-circle assay. (B) The schematic shows how newly formed telomeres at DR_L_-T2 were detected and sequenced. UDL6R-STELA was used to amplify telomeres at DR_L_-T2 and it occasionally amplified telomeres that were longer than the length of DR_L_-T2. Six of these intermediate length STELA products from three different DNA samples were re-amplified in a semi-nested, secondary PCR using primer DR8RT2 and Telorette2. Following gel extraction these products were Sanger sequenced using DR8RT2 or Telorette2. The TTAGGG repeats were visualized with FinchTV and counted manually. Electropherograms from one reamplified molecule from NWA008 are shown. As expected, the PAC2 motif was present (boxed sequence). A black line at base 273 shows where DR_L_-T2 in this sample was expected to end (after 18 TTAGGG repeats). Over 100 telomere repeats were counted from Telorette 2, considerably more than the length of T2 showing that this telomere has been lengthened. (C) Detecting telomerase activity in iciHHV-6B lymphoblastoid cell lines. Telomere repeat amplification protocol (TRAP) was used to detect low levels of telomerase activity in various iciHHV- 6B lymphoblastoid cell lines. Water and CHAPS were used as negative controls and cell lysate from a telomerase positive HUV-EC cell line was used as a positive control.

**Figure 6 – figure supplement 1.**
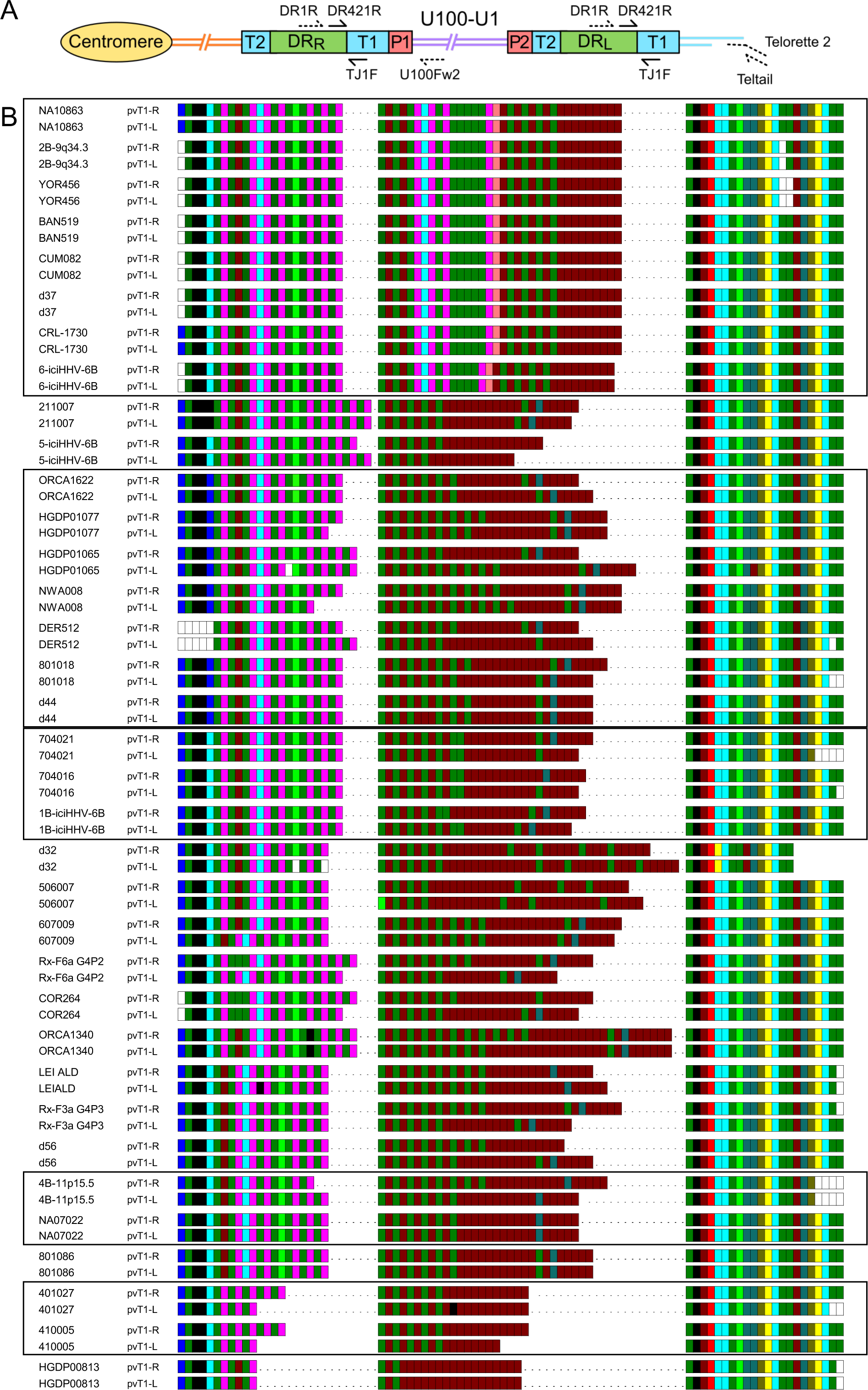
The majority of iciHHV-6B genomes show sequence differences between DR_R_-pvT1 and DR_L_-pvT1. DR_R_-T1 and DR_L_-T1 were specifically amplified by PCR using DR1R and U100Fw2 for DR_R_-T1, or by STELA using DR1R and STELA oligonucleotides (Teltail and Telorette 2) for DR_L_-T1. The products from DR_R_-T1 and DR_L_-T1 specific PCRs were used as input for secondary, nested PCR using DR421R and TJ1F to specifically amplify DR_R_-pvT1 and DR_L_-pvT1. Sanger sequencing was carried out using the TJ1F primer. Sequences were manually examined to generate pvT1 repeat maps. Repeats maps were colour coded and aligned as in Figure 1 – figure supplement 1. The majority of iciHHV-6B genomes (24/35, 68.6%) showed differences between DR_R_-pvT1 and DR_L_- pvT1, usually as loss or gain of a small number of hexameric telomere (TTAGGG) or degenerate telomere-like repeats. In a smaller number of cases, a single base change converted one repeat type to another. Regions containing CTAGGG repeats were most prone to loss or gain of repeats. Where differences between DR_R_-pvT1 and DR_L_-pvT1 were detected in multiple iciHHV-6B genomes from the same integration group (boxed) the differences were rarely the same between individuals. This indicates that the differences arose after integration. Interestingly, 9q iciHHV-6B genomes did not display any variation between DR_R_-pvT1 and DR_L_-pvT1.

**Supplementary Table 1.**
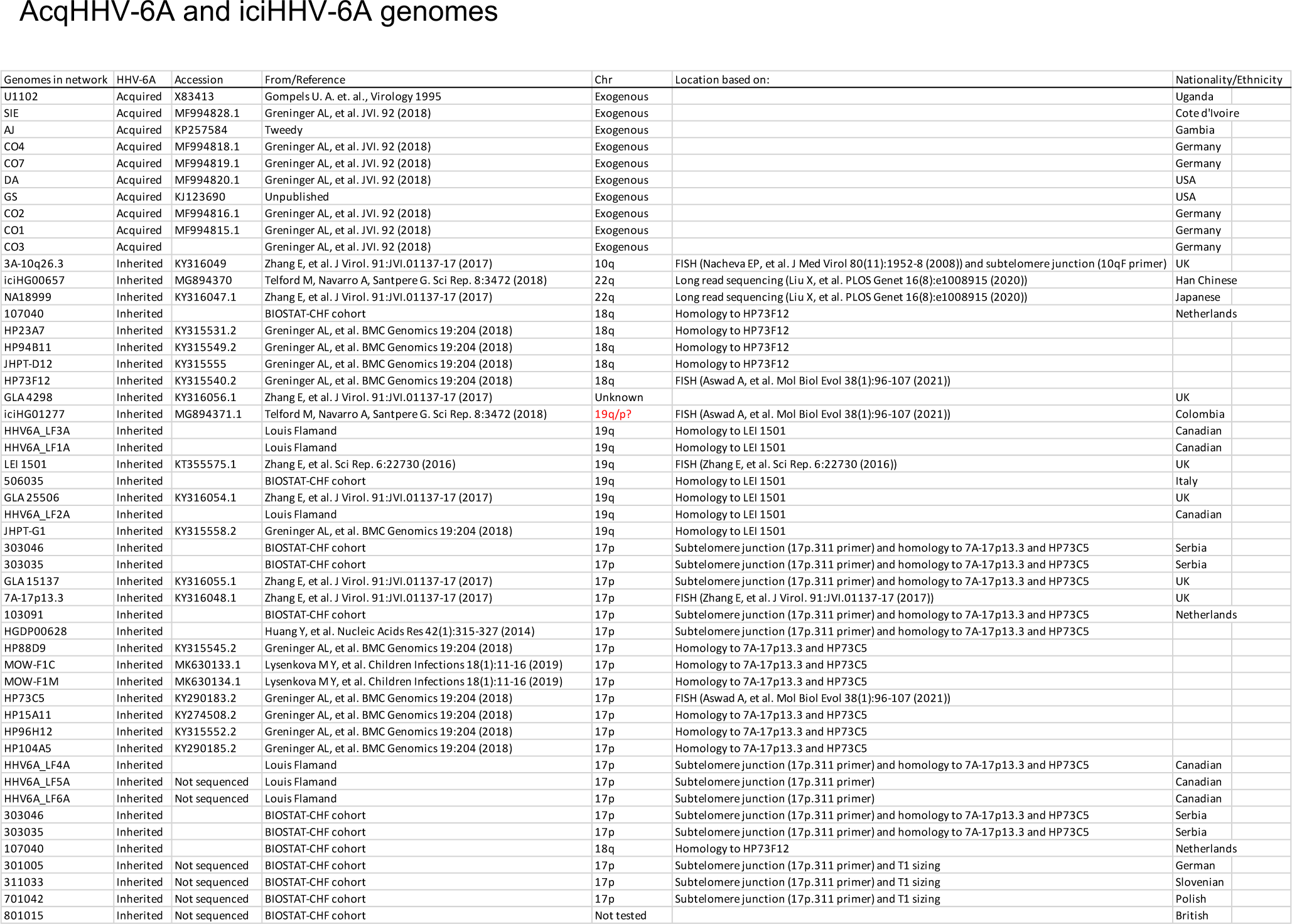
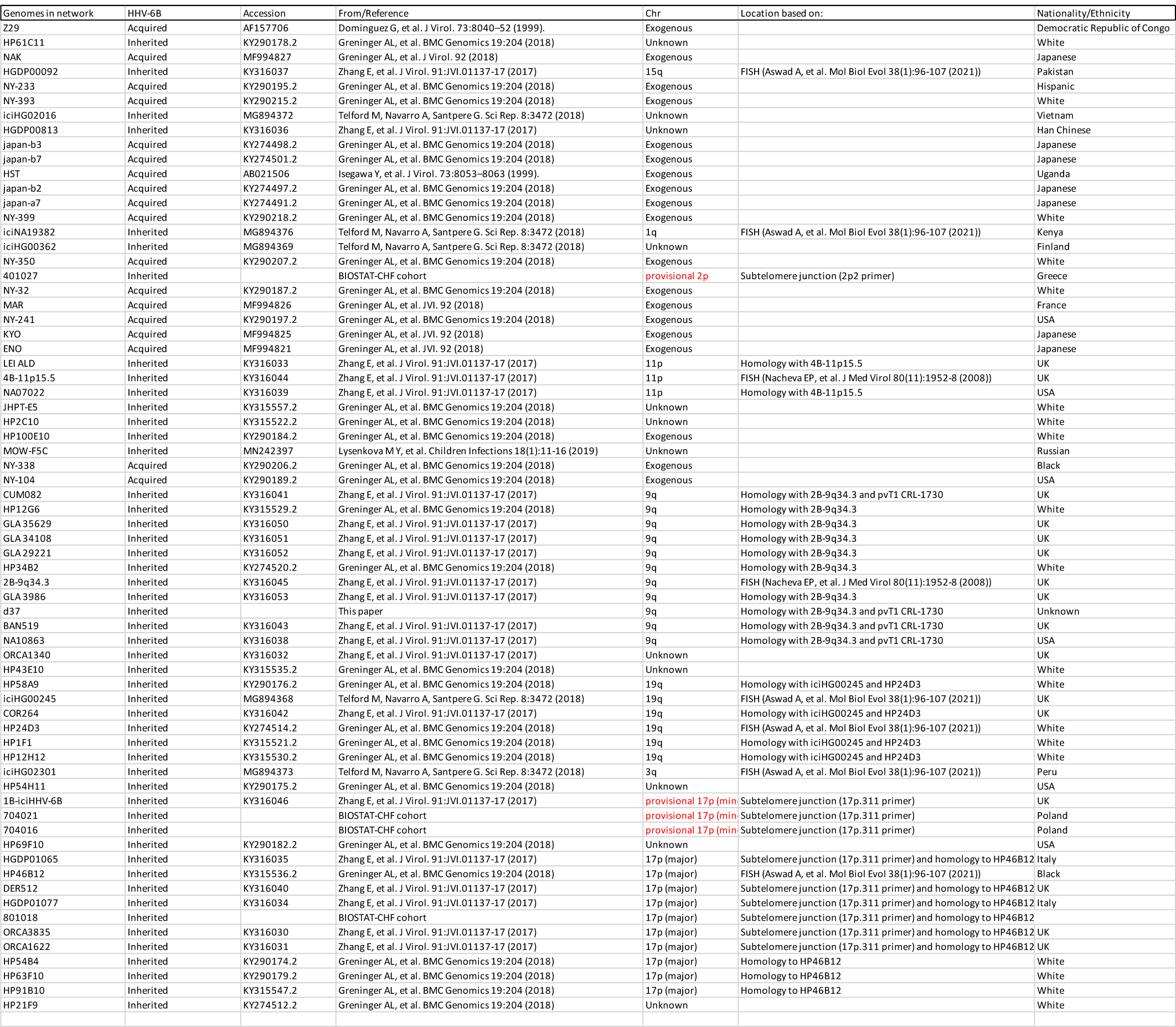
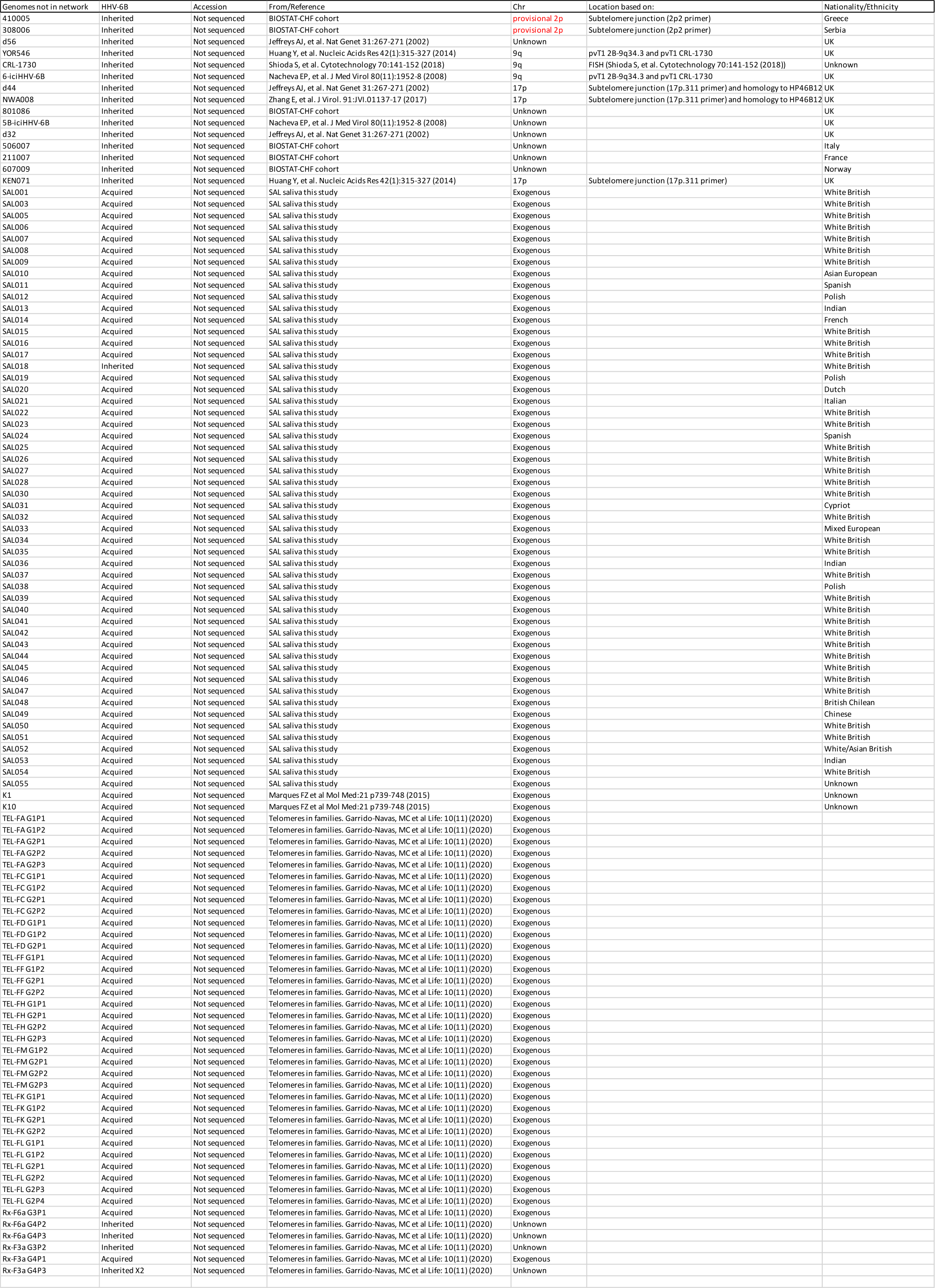
AcqHHV-6A/6B and iciHHV-6A/6B genomes included in study

**Supplementary Table 2.** Estimated Time to Most Recent Common Ancestor (TMRCA) for carriers of iciHHV-6A and iciHHV-6B with different chromosomal locations.

| Species | Chromosome location | No. genomes | TMRCA ± SD |
| --- | --- | --- | --- |
| HHV-6A | 17p | 14 | 105 033 ± 19 139 |
| HHV-6A | 18q | 5 | 22 976 ± 10 275 |
| HHV-6A | 19q | 8 | 68 928 ± 14 070 |
| HHV-6B | 9q | 11 | 20 855 ± 12 552 |
| HHV-6B | 17p (major group) | 7 | 23 409 ± 11 226 |
| HHV-6B | 19q | 6 | 24 579 ± 11 260 |

**Supplementary Table 3.** Measuring the percentage of acquired HHV-6B with a telomere, as an indicator of integration

|  | SAL015 | SAL027 | SAL039 | SAL040 | SAL044 | Tel-FA<br>G1P2 | K1 | K10 |
| --- | --- | --- | --- | --- | --- | --- | --- | --- |
| Total DNA input (ng) | 3600 | 1800 | 1800 | 900 | 900 | 1800 | 225 | 225 |
| Cell equivalent | 5.45 x 10<sup>5</sup> | 2.73 x 10<sup>5</sup> | 2.73 x 10<sup>5</sup> | 1.36 x 10<sup>5</sup> | 1.36 x 10<sup>5</sup> | 2.73 x 10<sup>5</sup> | 3.41 x 10<sup>4</sup> | 3.41 x 10<sup>4</sup> |
| Reactions with HHV-6B telomere | 5 | 3 | 9 | 2 | 19 | 4 | 0 | 4 |
| Copies of ciHHV-6 per cell | 9.17 x 10<sup>-6</sup> | 1.1 x 10<sup>-5</sup> | 3.30 x 10<sup>-5</sup> | 1.47 x 10<sup>-5</sup> | 1.39 x 10<sup>-4</sup> | 1.47 x 10<sup>-5</sup> | 0 | 1.17 x 10<sup>-4</sup> |
| Copies of HHV-6B per cell | 0.0027 | 0.002933 | 0.001667 | 0.001333 | 0.012467 | 0.00087 | 0.000525 | 0.0124 |
| Percentage integrated /% | 0.34 | 0.38 | 1.98 | 1.1 | 1.12 | 1.69 | 0 | 0.95 |

**Supplementary Table 4.** Variation between samples in the frequency of truncations at DR_L_-T2 and percentage lengthened.

| Sample name | DNA source | Truncations at DRL-T2 per cell | Percentage lengthened | Lengthened per cell |
| --- | --- | --- | --- | --- |
| NWA008 | Lymphoblasts | 0.0169 | 14.6 | 0.0025 |
| CEPH1375.02 | Lymphoblasts | 0.0091 | 4.55 | 0.0004 |
| COR264 | Lymphoblasts | 0.0078 | 15.8 | 0.0012 |
| 4B-11p15.5 | Lymphoblasts | 0.0248 | 3.33 | 0.0008 |
| 5B-17p13.3 | Lymphoblasts | 0.0158 | 0.00 | 0.00 |
| YOR546 | Lymphoblasts | 0.0136 | 18.2 | 0.0025 |
| OL | Lymphoblasts | 0.0172 | 0.00 | 0.00 |
| d37 | Pluripotent cells | 0.0093 | 80.9 | 0.0076 |
| CRL-1730 | Pluripotent cells | 0.019 | 37.0 | 0.007 |
| 401027 | Blood | 0.0025 | 63.7 | 0.0016 |
| 801018 | Blood | 0.0016 | 28.6 | 0.0005 |
| 211007 | Blood | 0.0025 | 77. 8 | 0.0019 |
| 704016 | Blood | 0.0009 | 100 | 0.0009 |
| 704021 | Blood | 0.0007 | 33.3 | 0.0002 |
| 801086 | Blood | 0.0032 | 68.8 | 0.0022 |
| 506007 | Blood | 0.0027 | 80.0 | 0.0022 |
| 607009 | Blood | 0.0084 | 67.6 | 0.0057 |
| 410005 | Blood | 0.0064 | 96.4 | 0.0061 |
| d32 | Sperm | 0.0063 | 4.41 | 0.0003 |
| d44 | Sperm | 0.0066 | 6.25 | 0.0004 |
| d56 | Sperm | 0.0059 | 8.87 | 0.0005 |
| Rx-F6a G4P2 | Saliva | 0.005 | 66.7 | 0.0033 |
| Rx-F6a G4P3 | Saliva | 0.0012 | 66.7 | 0.0008 |

**Supplementary Table 5.** Measuring the frequency of truncations at DR_R_-T1 in various samples

| Sample name | DNA source | No. Cells<br>(estimated from<br>total DNA<br>analysed) | No. of<br>STELA<br>reactions | No. of<br>STELA<br>products | No. of<br>reactions<br>with pvT1<sub>R</sub> | Estimated DR<sub>R</sub>-<br>T1 telomeres<br>per cell |
| --- | --- | --- | --- | --- | --- | --- |
| NWA008 | Lymphoblasts | 4848 | 64 | 172 | 1 | 0.0002 |
| COR264 | Lymphoblasts | 4848 | 64 | 123 | 7 | 0.0014 |
| HGDP01065 | Lymphoblasts | 3864 | 64 | 61 | 11 | 0.0028 |
| 401027 | Whole blood | 8712 | 98 | 393 | 8 | 0.0009 |
| 211007 | Whole blood | 4848 | 64 | 143 | 3 | 0.0006 |
| 704021 | Whole blood | 2424 | 32 | 201 | 3 | 0.0012 |
| 506007 | Whole blood | 2424 | 32 | 200 | 7 | 0.0029 |
| d32 | Sperm | 4848 | 64 | 65 | 6 | 0.0012 |
| d56 | Sperm | 2424 | 32 | 70 | 4 | 0.0017 |
| Rx-F6a G2P2 | Saliva | 4848 | 64 | 132 | 4 | 0.0008 |

**Supplementary Table 6.**
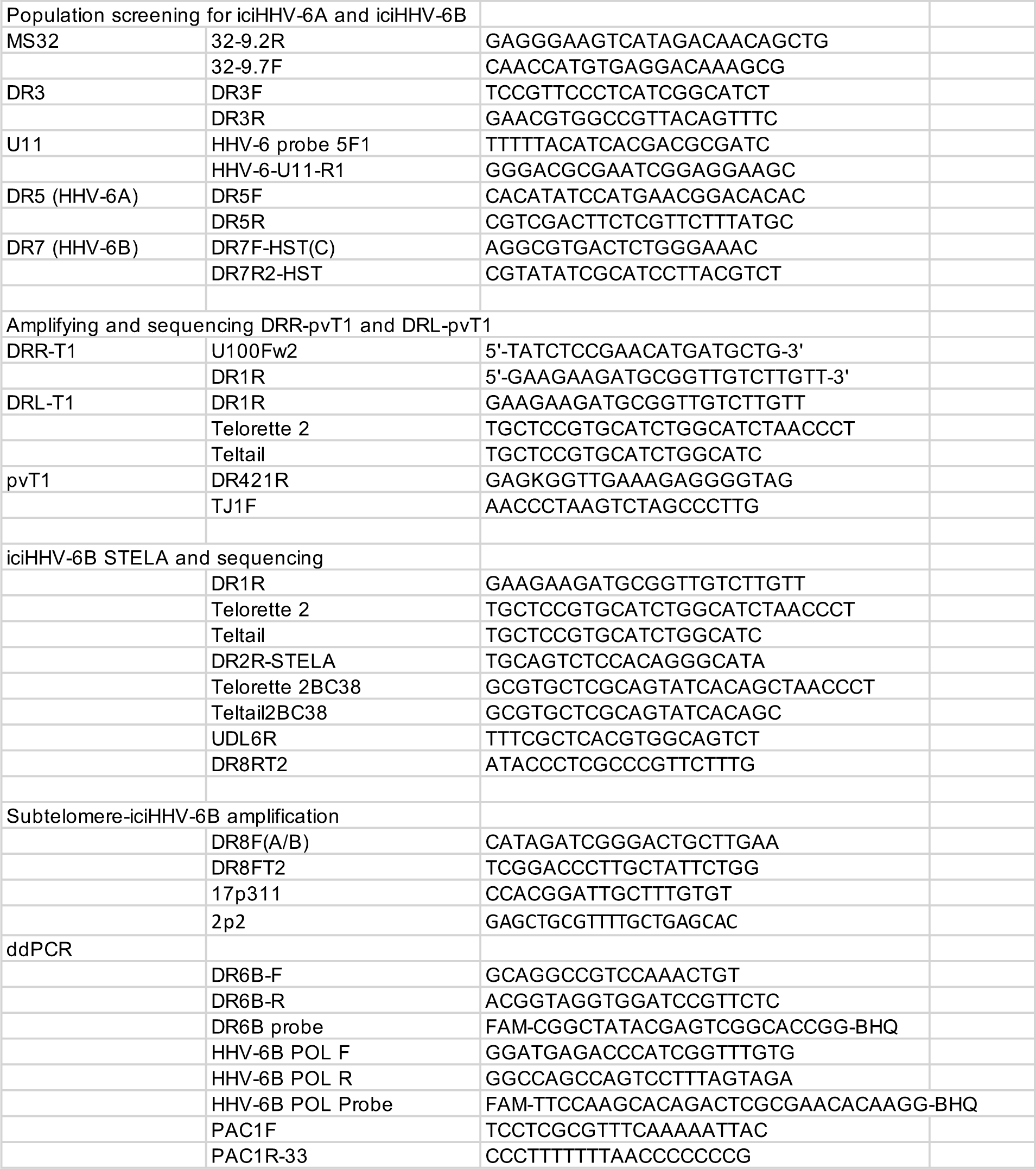
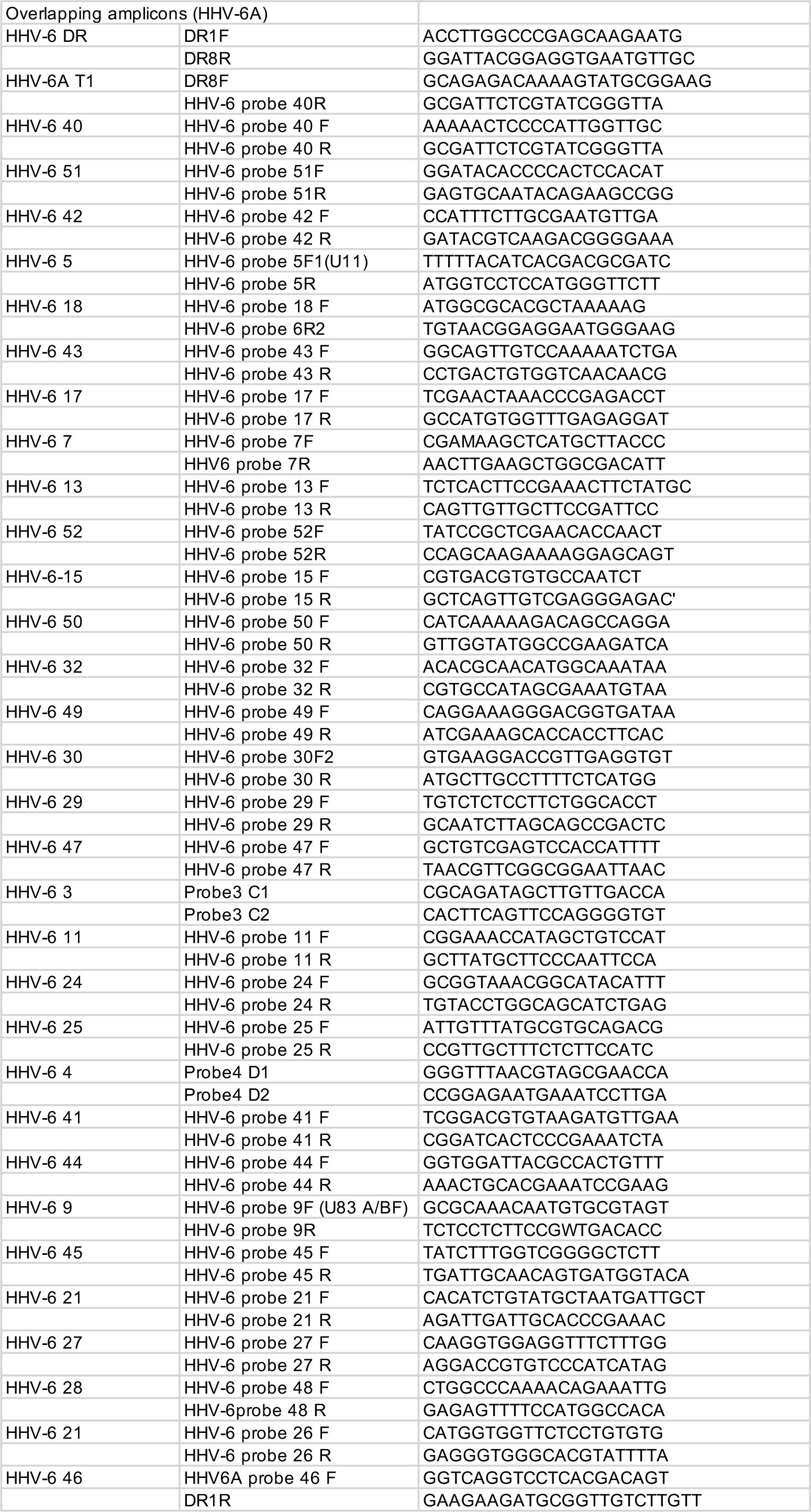
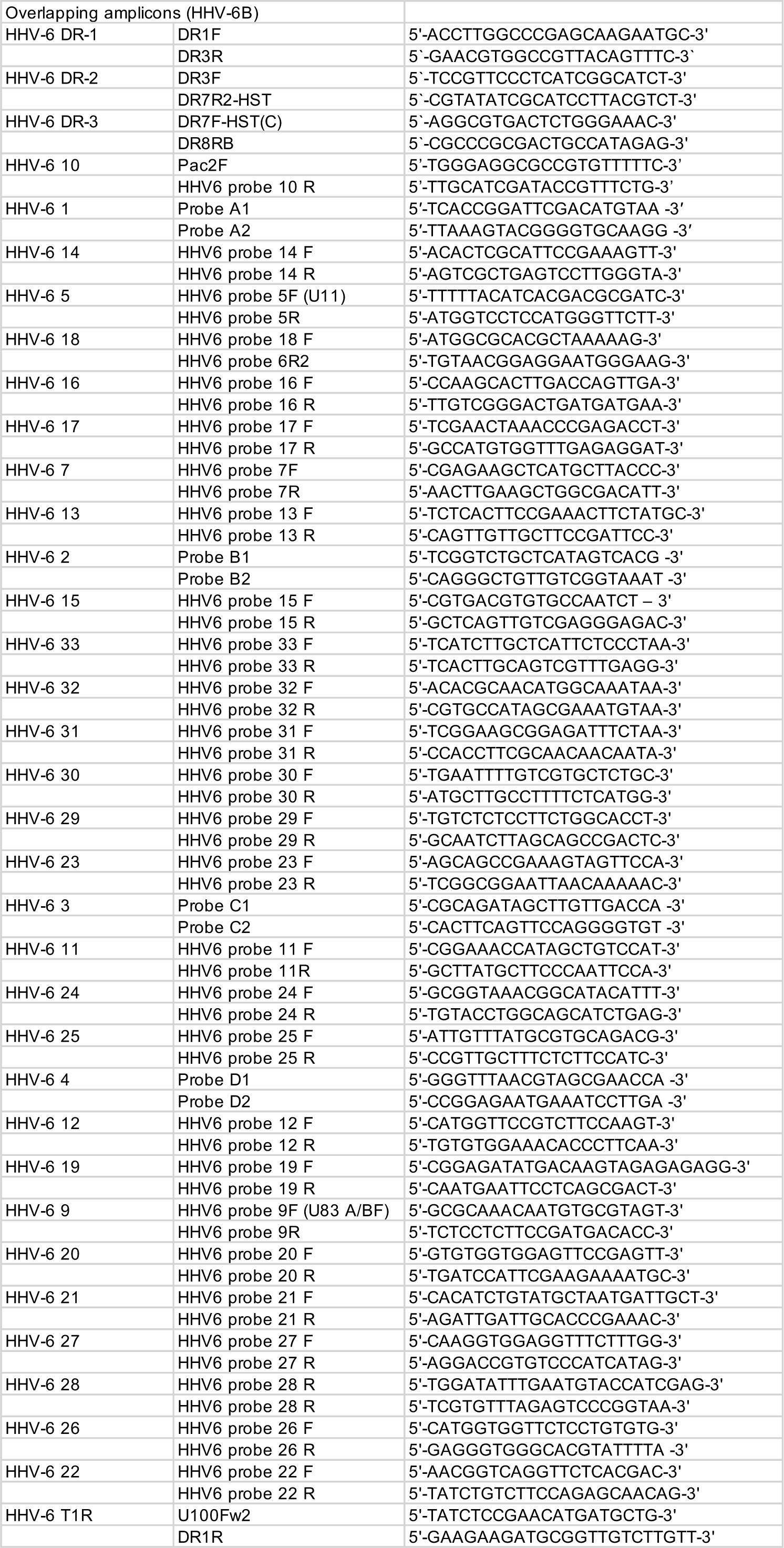
Primers used in this study

